# Secondary coordination sphere-controlled redox modulation drives the evolution of a metalloenzyme’s metal preference

**DOI:** 10.1101/2025.10.23.684290

**Authors:** ES Mackenzie, KM Sendra, A Baslé, R Mazgaj, TE Kehl-Fie, KJ Waldron

**Author notes:** Authors to whom correspondence should be addressed: Thomas E. Kehl-Fie,. Kevin J. Waldron,.

## Abstract

Changes in protein properties and functions are central to the evolution of life. Metalloproteins can evolve by changing their preference from one metal cofactor to another. Recently, we demonstrated that the widely distributed iron or manganese dependent superoxide dismutase (SodFM) family have undergone numerous metal-preference changes, including during evolutionary adaptation of pathogenic bacteria to altered metal availability within the host. Yet the underlying properties of metal-binding sites that control metalloenzyme metal-preference are unclear, and thus we lack an understanding of how enzymatic metal-preference can be re-shaped by evolution. Here, we used spectral features of bound iron or manganese, whose intensities reflect their oxidation state, to assess how their redox properties are manipulated during SodFM evolution. We systematically analysed the metal oxidation state across diverse SodFMs from multiple phylogenetic groups with different catalytic metal-preferences, including those known to have undergone evolutionary metal-preference switching. We observed a striking relationship between resting oxidation state and catalytic metal-preferences. Mutagenesis of second-sphere residues previously identified as determining metal preference revealed that they modulate metal-dependent activity and cofactor oxidation state in tandem, demonstrating these properties are linked. Together, these data argue that the differing SodFM metal preferences observed across the tree of life evolved through tuning of their redox properties by the secondary coordination sphere. This study gives insight into the process by which a metalloenzyme originally optimised for one metal cofactor can evolve a new metal preference, under suitable selection pressure, through re-optimisation of its active site for catalytic reactivity of the new metal cofactor.

**Significance statement:** Metal cofactors are needed by almost half of all enzymes. Catalytic metal-preference of metalloenzymes can evolve, for example to adapt to altered environmental metal availability. Yet, it is unclear how this evolutionary process occurs, enabling an active site optimised for one metal to change to become optimised for the new metal. Here, we have investigated this evolutionary process in a family of superoxide dismutase (SodFM) enzymes. We found that spectral features, which reflect the enzyme’s redox properties, of a diverse array of SodFMs with differing metal-preferences, and of mutated forms with artificially altered preferences, correlate with the metal-dependence of their activity. The data show that metal-preference change in SodFMs involves evolutionary tuning of the redox properties of SodFMs.

## Introduction

The ability of proteins to change their properties and functions is fundamental to the evolution of life. For example, evolutionary changes in metalloenzymes can enable them to utilise a new metal cofactor while retaining their ancestral activity, or can facilitate acquisition of entirely new catalytic functions^1–3^. Almost half of all enzymes utilise metal cofactors, reflecting their ancient and essential recruitment by biology^4–6^. Yet the evolutionary and biophysical mechanisms by which protein metal utilisation evolves remains to be fully elucidated.

We recently established the iron (Fe) or manganese (Mn) superoxide dismutase (SOD) enzyme family (SodFM) as a model system for studying metal preference evolution^2^. SODs utilise redox-active metal ions to catalyse detoxification of the reactive oxygen species superoxide, and play critical roles in human health, including the ability of microbes to survive within the host during infection^7,8^. SodFMs are the most prevalent SOD type across the tree of life^2^. Contrary to previous classification of SodFMs into three distinct sub-types according to their metal specificity^4,7,9,10^, we demonstrated their catalysis display a spectrum of metal-preference, ranging from strongly Mn-preferring, through metal-interchangeable (cambialistic), to strongly Fe-preferring^1,2^. Importantly, catalytic metal-preference is not constrained within SodFM phylogenetic groups^2^. Four of the five identified SodFM subfamilies (denoted SodFM1-5) contain isozymes with varying metal-preference, demonstrating their metal utilisation is evolutionarily dynamic^2^. Multiple independent SodFM metal preference changes have occurred, ranging from very ancient to very recent evolutionary modulations. Importantly, cambialistic SodFMs that exhibit activity with either cofactor have likely evolved from more metal-specific ancestors on multiple independent occasions. The most recent identified emergence of cambialism occurred during the proposed neofunctionalisation of a duplicated SodFM in *Staphylococcus aureus*^1^, which enabled this pathogenic bacterium to circumvent the Mn-deprivation it experiences during infection imposed by host nutritional immunity^8^. Thus, evolution can manipulate the metal-preference of SodFMs, and these changes can occur on relatively short evolutionary timescales^1,2^.

The cofactor flexibility of cambialistic SodFMs challenges the assumption that oxidoreductase metalloenzymes are highly specific for their cognate metal^11–13^. Their presence in diverse organisms, including pathogenic microbes, emphasises our limited understanding of metalloenzyme metal utilisation and the need to address this gap in knowledge. Crucially, despite their different metal-preferences, all SodFMs are related in sequence and share a conserved architecture, including the structure of their metal-binding active site^14–17^. Amino acids within the metal’s secondary coordination sphere, including a key residue denoted X_D-2_ (reflecting its position in the primary sequence relative to the conserved metal-coordinating aspartate), have frequently been involved in these evolutionary metal-preference modulation events^1,2^. However, it is unclear how such minor mutations can drive the biochemical changes observed during SodFM evolution.

Although other models have been proposed^18–20^, a ‘redox tuning’ model has gained some limited empirical support as an explanation of the differences in metal-preference between the *E. coli* MnSOD and FeSOD^21^. This model hypothesises that these SodFM isozymes differentially manipulate the intrinsic reduction potentials of Fe^2+^/Fe^3+^ (*E*^0^ = 0.77 V) and Mn^2+^/Mn^3+^ (*E*^0^ = 1.51 V) to be close to the optimal potential for catalysing both steps of the chemical reaction (*E*^0^ ≈ 0.3 V) with its cognate ion, with MnSOD having to apply a greater adjustment than FeSOD to achieve this optimum^21,22^. The resulting mismatch of tuning could explain why each of these SodFM isozymes have high turnover with one metal but are catalytically ineffective with the other metal, as the ‘wrong’ metal’s reduction potential will be inappropriate within an architecture optimised for the ‘correct’ metal^21,23^. Specifically, the MnSOD architecture would ‘over-tune’ the potential when loaded with Fe, and the FeSOD architecture would ‘under-tune’ bound Mn. The model also predicts that cambialistic SodFMs should be sub-optimally tuned but within the target range for both metals, and therefore should exhibit lower turnover than metal-preferring SodFMs. This would suggest that cambialism is an evolutionary trade-off, compromising higher turnover in favour of physiological flexibility from relaxed cofactor preference. When a SodFM evolves to utilise the alternative metal, thereby switching its metal preference, such a model would require evolutionary re-tuning of the reduction potential.

Direct testing of the redox tuning model is challenging because quantitative measurement of SodFM reduction potentials is not facile with existing methods^24–26^. To overcome these limitations, here we leveraged the spectral features of their bound *d*-block metals, Fe and Mn, whose intensities reflect their oxidation state, to assess the distribution of redox states of the population of metal ions in SodFM samples. We performed a systematic, semi-quantitative analysis of cofactor oxidation state under standardised assay conditions across a vast set of purified samples of diverse SodFM isozymes, from multiple phylogenetic groups and with a range of catalytic metal-preferences. Our sampling included SodFMs known to have undergone evolutionary metal-preference switching and a set of mutated variants whose preference had been artificially altered. We observed a striking trend in their redox state that followed their catalytic metal preferences, consistent with tuning of the metal’s redox properties by the protein architecture being an underlying biophysical principle of SodFM metal-preference evolution.

## Results

### Spectral properties of SodFMs at aerobic equilibrium reflect their metal preference

A set of seven bacterial SodFMs were initially selected for characterisation from the two main subfamilies, SodFM1 and SodFM2. We previously identified evolutionary metal-preference modulation events in both of these subfamilies^2^. Each enzyme was purified to homogeneity in both Fe- and Mn-loaded forms, in triplicate, and the metal preference of its enzymatic activity was quantitatively determined. A cambialism ratio (CR), defined as the Fe-dependent activity divided by the Mn-dependent activity^1^ (Table 1), was experimentally verified for each enzyme. This ratio approaches 0 for increasingly Mn-preferring SodFMs, has values close to 1 for cambialistic enzymes capable of utilising either metal with similar efficiency, and increases >2 with increasing Fe-preferring activity. All CR values were calculated for metal-verified protein preparations and corroborated the previously measured approximate CR values estimated using our standardised 24-well plate format activity assay in soluble extracts of *E. coli* cells expressing the heterologous SodFM isozymes^2^.

**Table 1.** Summary of CR and nOS data of all wild type SodFMs and mutant variants under study. Table summarising the key data collected for all wild type and mutated variants of the SodFMs under study herein. Data demonstrate each isozyme’s metal-preference (CR) and the determined redox state (nOS) as calculated from the UV/visible absorption spectra. Data here are a summary of those detailed in Source Data Table 1, which includes all individual results, errors and statistical analyses. ^a^Cambialism ratio (CR) values were calculated by dividing the measured Fe-dependent activity by the measured Mn-dependent activity, as determined by a NBT/riboflavin-based spectrophotometric assay. ^b^Normalised oxidation state (nOS) values were calculated by normalising the ‘resting’ UV-visible absorption spectral peak intensity (^Mn^λ_480nm_ and ^Fe^λ_350nm_) between the fully reduced peak intensity (nOS=0) and fully oxidised peak intensity (nOS=1) (Fig. 1). ^c^Only one fully metal-loaded form of *Ec*SodA (in both its wild type and mutated form) and *Ba*SodA2 could be produced, precluding measurement of activity of the other form, so CR values could not be calculated for these two isozymes. ^d^While all other SodFMs were analysed in triplicate, *Bf*Sod, *Am*Sod, *Cw*Sod and their mutant variants were analysed as supplementary samples with *n* = 1 in order to test whether the trend, established from the core *n* = 3 dataset, was also observed in these more highly evolutionarily divergent isoymes. ND=Not determined in this study.

| Species | Enzyme | Wild type |  |  |  | X<sub>D-2</sub>-X<sub>D-1</sub> mutant |  |  |  |
| --- | --- | --- | --- | --- | --- | --- | --- | --- | --- |
|  |  | SodFM | CR<sup>a</sup> | nOS<sub>Fe</sub><sup>b</sup> | nOS<sub>Mn</sub><sup>b</sup> | Mutation | CR<sup>a</sup> | nOS<sub>Fe</sub><sup>b</sup> | nOS<sub>Mn</sub><sup>b</sup> |
| *E. coli* | *EcSodA*<sup>c</sup> | FM1 | ND<sup>c</sup> | ND<sup>c</sup> | 0.64 | G<sub>D-2</sub>T-L<sub>D-1</sub>V<sup>c</sup> | ND<sup>c</sup> | ND<sup>c</sup> | 0.47 |
| *S. aureus* | *SaSodA* | FM1 | 0.09 | 0.06 | 0.53 | G<sub>D-2</sub>L-L<sub>D-1</sub>F | 0.49 | 0.16 | 0.27 |
| *B. subtilis* | *BsSodA* | FM1 | 0.13 | 0.09 | 0.56 | G<sub>D-2</sub>V-L<sub>D-1</sub>I | 1.49 | 0.23 | 0.24 |
| *B. anthracis* | *BaSodA1* | FM1 | 0.20 | 0.01 | 0.62 | G<sub>D-2</sub>V-L<sub>D-1</sub>I | 0.74 | 0.21 | 0.21 |
| *L. monocytogenes* | *LmSodA* | FM1 | 0.30 | 0.02 | 0.55 | G<sub>D-2</sub>V-L<sub>D-1</sub>I | 1.01 | 0.27 | 0.21 |
| *S. pyogenes* | *SpSodA* | FM1 | 0.38 | 0.16 | 0.58 | A<sub>D-2</sub>V-L<sub>D-1</sub>I | 0.99 | 0.37 | 0.39 |
| *B. fragilis* | *BfSodFM2*<sup>d</sup> | FM2 | 0.73 | 0.93<sup>d</sup> | 0.27<sup>d</sup> | G<sub>D-2</sub>T-F<sub>D-1</sub>C<sup>d</sup> | 24.7 | 0.86<sup>d</sup> | 0.01<sup>d</sup> |
| *S. aureus* | *SaSodM* | FM1 | 1.23 | 0.44 | 0.28 | L<sub>D-2</sub>G-F<sub>D-1</sub>L | 0.44 | 0.12 | 0.47 |
| *A. muciniphila* | *AmSodFM3*<sup>d</sup> | FM3 | 4.03 | 0.69<sup>d</sup> | 0.09<sup>d</sup> |  |  |  |  |
| *c. Wolfebacteria* | *cWSodFM1*<sup>d</sup> | FM1 | 18.5 | 0.47<sup>d</sup> | 0.00<sup>d</sup> | V<sub>D-2</sub>G<sup>d</sup> | 5.43 | 0.12<sup>d</sup> | 0.00<sup>d</sup> |
| *E. coli* | *EcSodB* | FM2 | 27.5 | 0.83 | 0.03 | T<sub>D-2</sub>G-V<sub>D-1</sub>L | 3.99 | 0.78 | 0.02 |
| *N. gonorrhoeae* | *NgSodB* | FM2 | 29.3 | 0.83 | 0.18 | T<sub>D-2</sub>G | 7.78 | 0.78 | 0.16 |
| *B. anthracis* | *BaSodA2*<sup>c</sup> | FM1 | ND<sup>c</sup> | 0.35 | ND<sup>c</sup> |  |  |  |  |

Quantitative measurement of SodFM reduction potentials is challenging. The buried active site is only accessible *via* a narrow solvent channel, necessitating the use of small-molecule electrochemical mediators to communicate between the active site metal and an electrode, which is slow and inefficient and results in protein degradation over the course of experiments^24–26^. Given these limitations, it is not experimentally possible to measure reduction potentials of a diverse array of SodFMs and their mutated variants^2^ to explicitly test a role for redox tuning in SodFM evolution. Here, we leveraged the spectral features of their *d*-block metals, Fe and Mn, to assess the redox properties of SodFMs (Fig. 1). Both metals exhibit absorptions in the UV-visible region of the spectrum^27,28^. Mn^3+^ yields a broad, distinct feature from 400-700 nm with a maximum at 480 nm (defined herein ^Mn^λ_480nm_), whereas Fe^3+^ gives rise to a shoulder on the main polypeptide absorbance with a maximum at 350 nm (herein ^Fe^λ_350nm_). These features give rise to visible purple (Mn) or brown (Fe) colours in concentrated SodFM samples. Crucially, the intensities of these colours and the spectral features were increased after complete oxidation to Mn^3+^/Fe^3+^ and diminished on complete reduction to the Mn^2+^/Fe^2+^ oxidation state (Fig. 1). This indicates that the resting state, representing samples at aerobic equilibrium under standardised experimental conditions, contained mixtures of reduced and oxidised protein-bound metal cofactors. We thus utilised the peak intensity of these absorption features to assess the resting oxidation state of the active site metals in SodFM samples.

**Fig. 1.**
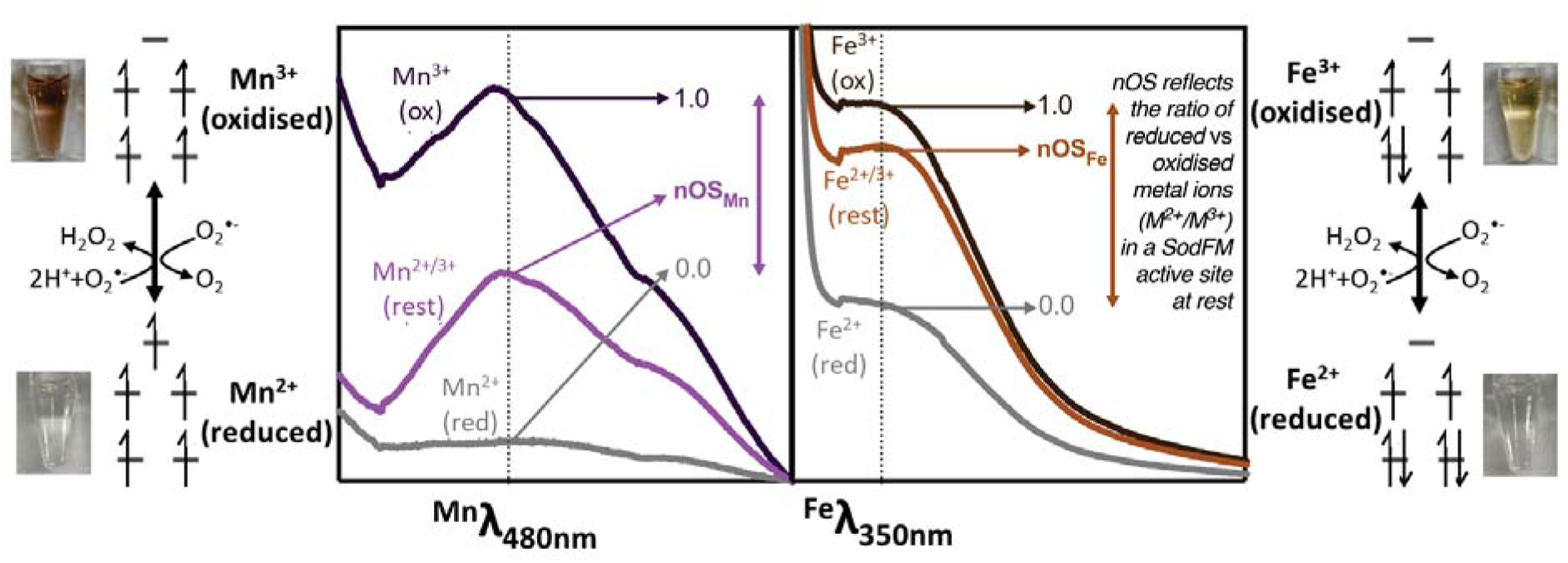
Schematic illustration of SodFM reaction, colour and spectra, and calculation of nOS. UV-visible absorption spectra (central panels) of *Sa*SodA (left) and *Ec*SodB (right) loaded with Mn and Fe, respectively. The spectra of each protein as purified, which were at equilibrium under atmospheric aerobic conditions, are termed ‘resting spectra’ and labelled M^2+^/M^3+^ (rest) and are shown with the lighter coloured lines (Mn-*Sa*SodA, purple; Fe-*Ec*SodB, brown). Spectra of the oxidised protein samples are labelled M^3+^ (ox) and are shown in darker coloured lines, and spectra of the reduced proteins are labelled M^2+^ (red) and are shown in grey. Spectra were obtained from samples containing approximately 100 μM protein in 20 mM Tris pH 7.5, 150 mM NaCl. Chemical oxidation was achieved by incubation with one mole equivalent of potassium permanganate and chemical reduction by incubation with three mole equivalents sodium dithionite for 10 min, followed by extensive buffer exchange using centrifugal filtration into the same buffer. The derivation of nOS values by normalisation of the resting spectral intensity, ^Mn^λ_480nm_ and ^Fe^λ_350nm_, against those of the fully oxidised and fully reduced samples is illustrated. Also shown (outer panels) are photographs of each oxidised (upper photo) and reduced (lower photo) protein samples, with their respective electronic configurations (high spin ions in a pseudo-trigonal bipyramidal configuration) and the superoxide dismutation reaction are schematically illustrated.

Initially, we sought to verify the relationship between the intensity of these spectral features and the metal’s oxidation state using three well-studied model SodFMs that represent archetypes of the three traditional ‘metal-specificities’: MnSOD (*S. aureus* SodA^8^, belonging to subfamily SodFM1), camSOD (*S. aureus* SodM^8^; SodFM1), and FeSOD (*E. coli* SodB^29^; SodFM2). We acquired resting, fully oxidised and fully reduced absorption spectra for each SodFM in both their Mn- and Fe-loaded forms under standardised experimental conditions (Fig. 2A). The resting ^Mn^λ_480nm_ was more intense for *Sa*SodA than for *Sa*SodM. This suggests that Mn-*Sa*SodA contained a higher proportion of oxidised Mn^3+^ cofactors. Conversely, the resting spectrum of catalytically inactive Mn-*Ec*SodB closely resembled that of its fully reduced form (Fig. 2A), resulting in ^Mn^λ_480nm_⁓0, suggesting it contained almost exclusively reduced Mn^2+^ cofactors. The reverse was true for the Fe-loaded forms (^Fe^λ_350nm_: *Ec*SodB>*Sa*SodM>*Sa*SodA; Fig. 2A). We noted that the spectral intensities followed the trend in their metal-dependent catalytic activities (Table 1); those of enzymatically active forms were consistent with mixed oxidation state populations at rest, whereas inactive forms were more extensively reduced.

**Fig. 2.**
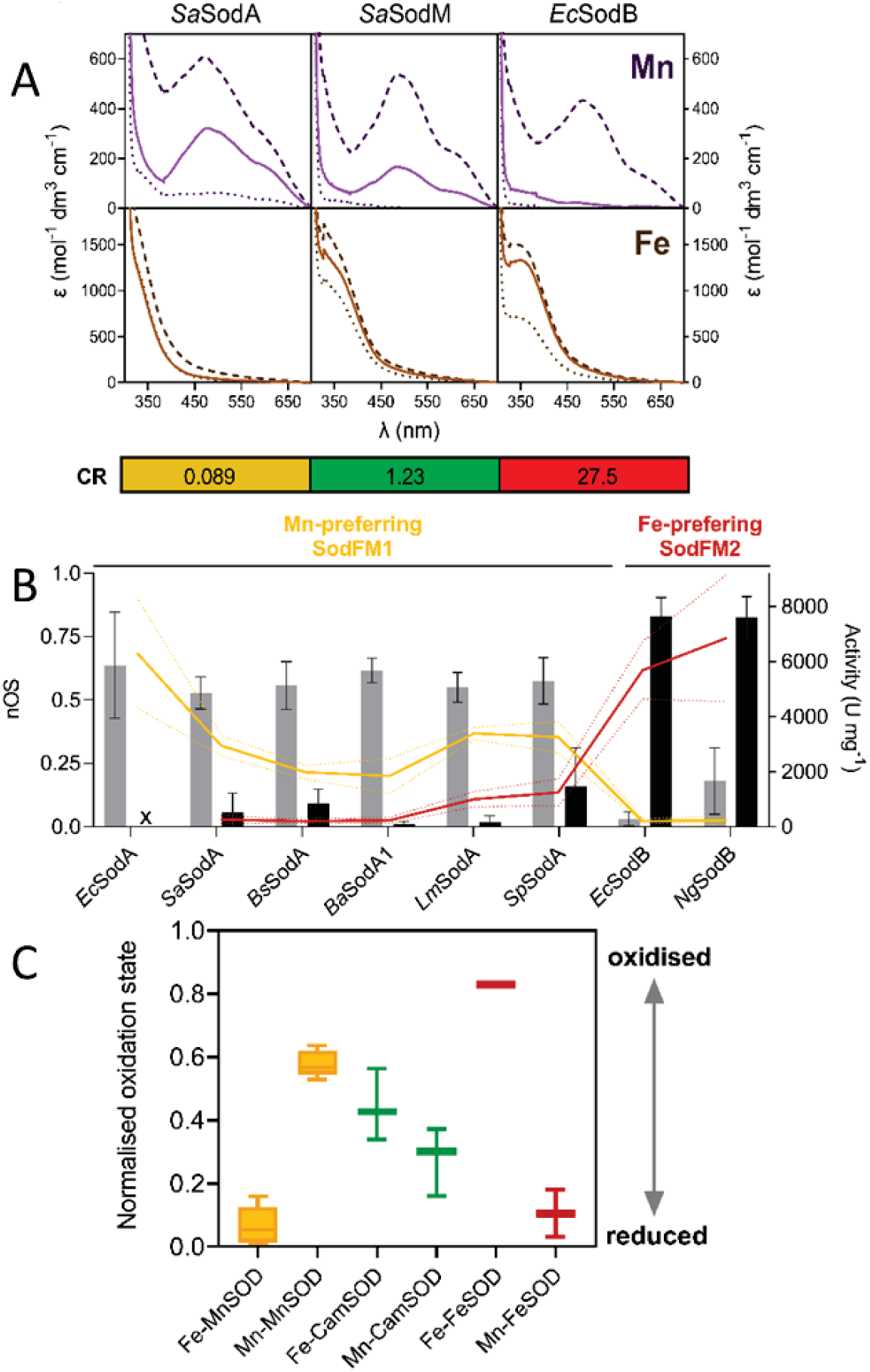
Spectra of SodFMs at rest correlate with their metal-dependent activity. **A** UV-visible absorption spectra of Mn-loaded (upper panels, purple lines) and Fe-loaded (lower panels, brown lines) *Sa*SodA, *Sa*SodM and *Ec*SodB. Resting spectra of proteins equilibrated under atmospheric conditions are shown as solid lines, and spectra of oxidised and reduced proteins as dashed and dotted lines, respectively. The resting spectra lie between the respective oxidised and reduced spectra for catalytically active forms (Mn-*Sa*SodA, Fe-*Ec*SodB, Mn-*Sa*SodM and Fe-*Sa*SodM) but are similar to reduced spectra for inactive forms (Fe-*Sa*SodA, Mn-*Ec*SodB). Spectra shown are *n*=1 but representative of further replicates (*n*=3). Cambialism ratio (CR) values are annotated below, coloured by Mn-preference (yellow), cambialistic (green), and Fe-preference (red). **B** Inverse trend between Mn-dependent (yellow line) and Fe-dependent activity (red line) detected for SodFM1 and SodFM2 isozymes and their normalised oxidation state (nOS) when loaded with Mn (grey bars) or Fe (black bars). The nOS values were calculated by normalising resting spectral peak intensity (^Mn^λ_480nm_ and ^Fe^λ_350nm_) between those of reduced (nOS=0) and oxidised (nOS=1) peak intensity. Enzymes were ordered along the x-axis according to their CR. All spectra and activities were measured in triplicate using independent biological replicates. Errors represent the standard deviation around the mean, displayed as error bars for nOS and dotted lines for activity in their respective colours. As we were unable to generate Fe-*Ec*SodA, no activity or nOS are displayed for this form (x indicates lack of data). **C** Box and whisker plot comparing nOS from triplicate analyses of each SodFM metal form. Whiskers were calculated using Tukey’s test (*n*=3).

The absolute spectra varied between these three control SodFM isozymes, so peak spectral intensities were normalised to enable semi-quantitative comparisons. Taking the spectra of the fully reduced and fully oxidised samples as representing the lower and upper bounds of absorbance, respectively, we generated normalised numerical values for ^Mn^λ_480nm_ and ^Fe^λ_350nm_ from each enzyme’s resting spectrum (Fig. 1). The resulting values are taken to approximate the proportion of active site metal ions in the resting sample that are oxidised, and are therefore termed normalised oxidation state (nOS), because nOS=0 represents fully reduced and nOS=1 represents fully oxidised samples. We found that all catalytically active forms of these SodFMs exhibited nOS values >0.25, indicating a significant proportion of oxidised metal cofactors at rest (Table 1). These nOS values are consistent with the ability to perform both the oxidative and reductive half-reactions of the SOD catalytic cycle under these conditions as the enzyme population at rest contains cofactors poised to perform either half-reaction. Conversely, the two SOD forms lacking activity, Fe-*Sa*SodA and Mn-*Ec*SodB, had nOS values close to zero, implying these samples contained exclusively reduced cofactors at rest (Table 1).

To validate these results, a larger panel of purified isozymes from the SodFM1 and SodFM2 subfamilies^2^, including enzymes from *Bacillus subtilis*, *Listeria monocytogenes*, *Streptococcus pyogenes* and *Neisseria gonorrhoeae*, was analysed. The spectra of these Mn-preferring SodFM1 (*Bs*SodA, *Lm*SodA and *Sp*SodA) and Fe-preferring SodFM2 (*Ng*SodB) isozymes^2^ (Supp. Fig. S1) were consistent with those of the canonical enzymes from *S. aureus* and *E. coli* (Fig. 2B). All the SodFM1s exhibited higher nOS_Mn_ values (0.53–0.64) but lower nOS_Fe_ (0.02–0.16), consistent with the strong Mn-preference of their catalysis. The SodFM2 isozyme had a higher nOS_Fe_ (0.83) and a low nOS_Mn_ (0.03), consistent with its strong Fe-preference (Table 1).

Taken together, the data indicated that in SodFM1 and SodFM2, the two subfamilies most commonly found across the tree of life^2^, the enzyme’s nOS values reflected their metal preference (Fig. 2C). When loaded with the wrong metal, the highly metal-preferring SODs had few oxidised cofactors at rest (nOS≈0), but were much more highly oxidised when loaded with their preferred cofactors. Notably, the cambialistic SOD had higher nOS values with both metals reflecting the presence of oxidised cofactors in both forms. These data suggested a trend between resting oxidation state and catalytic metal-preference. Crucially, this trend was conserved across the SodFM1 and SodFM2 subfamilies that are derived from an ancient evolutionary split^2^, suggesting the hypothesis that evolutionary processes can shift catalytic metal-preference by altering this biophysical property.

### Both metal preference and nOS are regulated by the SodFM secondary coordination sphere

Numerous studies have demonstrated the role in determining SodFM metal-preference played by residues localised within the metal cofactor’s secondary coordination sphere^1,2,18,20,22,30–32^. These residues have been altered during evolutionary adaptations that have resulted in changes of metal-preference of SodFM catalysis^1,2^. We hypothesised that metal-preference changes caused by mutations in these residues would also be associated with changes in nOS. To test this, we calculated nOS values for a set of SodFM variants whose metal-preference had been altered through mutation of second sphere residues (Fig. 3A). Mutations were designed to mimic observed evolutionary events of metal preference modulation^2^, by switching native residues for those observed in the same locus in other SodFM isozymes that have contrasting metal preferences.

**Fig. 3.**
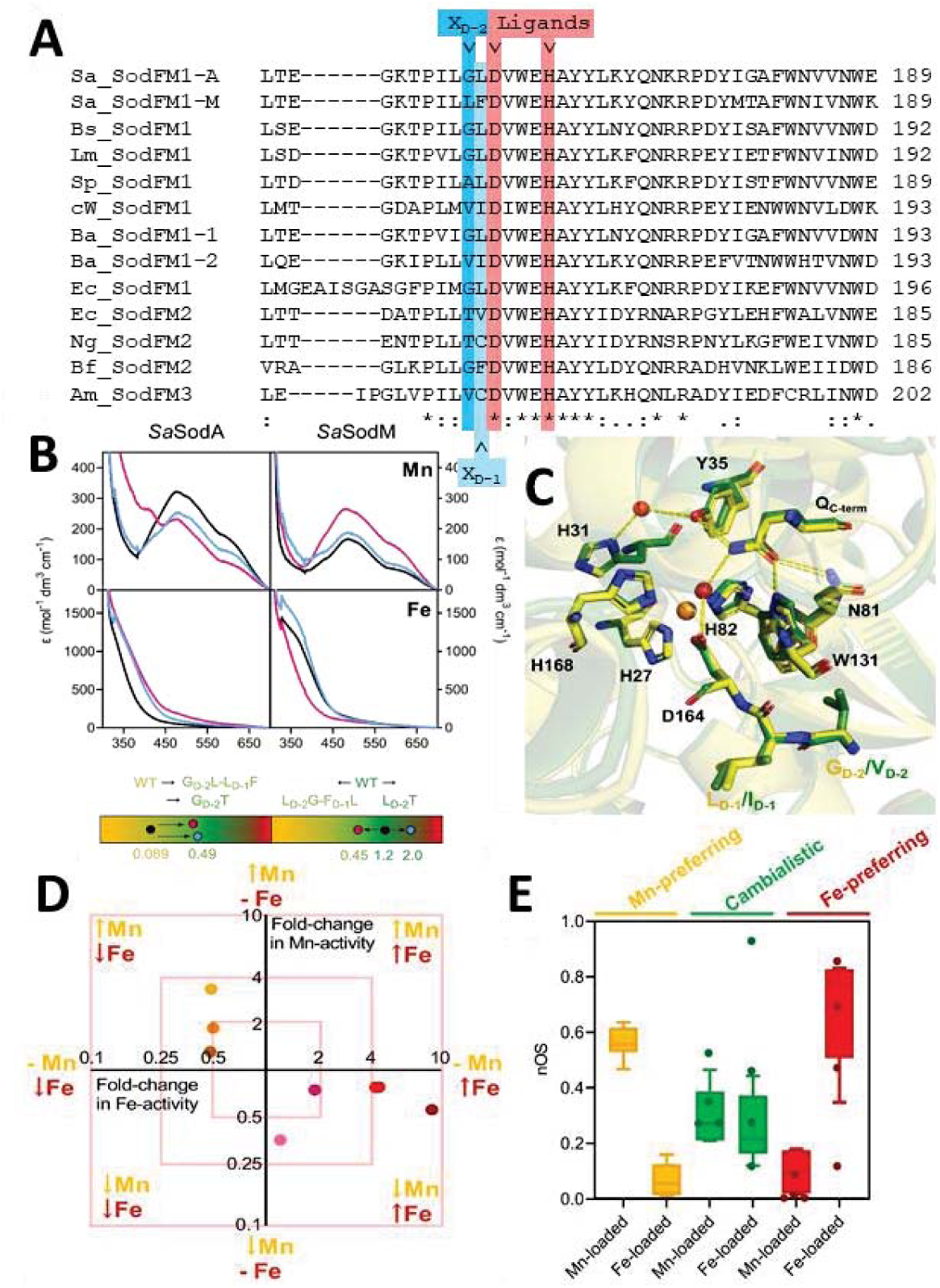
Spectra and nOS in mutated SodFM variants with altered metal preference. **A** Section of a sequence alignment of all SodFMs in this study (Sa=*Staphylococcus aureus*; Bs=*Bacillus subtilis*; Lm=*Listeria monocytogenes*; Sp=*Streptococcus pyogenes*; cW=candidatus *Wolfebacteria*; Ba=*Bacillus anthracis*; Ec=*Escherichia coli*; Ng=*Neisseria gonorrhoeae*; Bf=*Bacteroides fragilis*; Am=*Akkermansia muciniphila*). Second sphere residues targeted for mutagenesis, X_D-2_ (dark blue) and X_D-1_ (light blue), and two of the conserved metal ligands (red) are highlighted. Alignment was produced with Clustal Omega, and conservation levels are shown by symbols below the alignment where ‘*’ represents fully conserved residues, ‘:’ represents high and ‘.’ represents low level of conservation. **B** UV-visible absorption resting spectra of *Sa*SodA (left) and *Sa*SodM (right) when Mn- (upper panels) and Fe-loaded (lower panels) for wild type (black), X_D-2_/X_D-1_ (pink) and X_D-2_T (blue) mutant variants. Tricolour gradient panels below show the shift in CR from wild type (black circles) to X_D-2_/X_D-1_ (pink circles) and X_D-2_T (blue circles) variants. Changes in spectral intensity were seen where mutagenesis caused a shift in metal preference. **C** Ribbon and stick representation of superimposed (RMSD=0.38 Å) active sites of *Lm*SodA (yellow; 1.65 Å) and the *Lm*SodA VD-2-I_D-1_ variant (green; 1.40 Å) displaying metal-coordinating ligands, water-coordinating glutamine, and X_D-2_X_D-1_ residues. **D** Plot shows fold changes (log2 scale) in activity with Mn (orange) and Fe (red) of tested SodFM mutants relative to wild types (Table 1). Variants either gained Fe activity and lost Mn activity (*Sa*SodA, red; *Bs*SodA burgundy, *Ba*SodA1 dark red; *Lm*SodA, light pink; *Sp*SodA, dark pink), or gained Mn activity and lost Fe activity (*Sa*SodM, brown; *Ec*SodB, yellow; *Ng*SodB, orange). **E** Box and whisker plot of nOS from replicated wild type and mutant SodFM analyses, grouped by metal preference. Whiskers calculated using Tukey’s test (*n* = 3), nOS were calculated from spectra as previously described. Overlaid points display nOS of divergent wild type and mutant SodFMs analysed in single replicate (*n* = 1).

We previously demonstrated the residue X_D-2_ was key to the evolutionary divergence of the staphylococcal SodFM1s^1^. Its neighbouring residue, X_D-1_, plays a moderating role in metal-preference^1^. X_D-2_ was also shown to be important in the evolution of metal-preference in other SodFMs sampled across the tree of life^2^. Reciprocal mutagenesis of this amino acid pair in the *S. aureus* SodFMs (G_D-2_-L_D-1_ in Mn-preferring *Sa*SodA, CR=0.089; L_D-2_-F_D-1_ in cambialistic *Sa*SodM, CR=1.225) substantially inverted their metal preferences as well as inverting the susceptibility of their Mn^3+^ forms to chemical reduction by dithionite^1^, consistent with these residues modulating redox properties of the metal. The *S. aureus* SodFMs are a model of metal-preference evolution because both their physiological role during infection^8^ and their biochemical and structural properties^1,33^ were previously established. Here, we first confirmed that mutagenesis of X_D-2_/X_D-1_ created a more cambialistic variant of *Sa*SodA (L_D-2_-F_D-1_, CR=0.490) and a more Mn-preferring variant of *Sa*SodM (G_D-2_-L_D-1_, CR=0.448) (Table 1). Variants of both *Sa*SODs were also created with the non-native substitution T_D-2_, a residue common in isozymes of subfamily SodFM2^2^ that is most commonly indicative of Fe-preferring SODs^2,9,22,23^. Indeed, this mutation increased Fe-dependent activity in both cases (*Sa*SodA T_D-2_, CR=0.594; *Sa*SodM T_D-2_, CR=1.991) (Table 1). Crucially, all of these mutations also affected the resting spectra (Fig. 3B) and the resulting nOS values (Table 1). The variants *Sa*SodA-L_D-2_-F_D-1_, *Sa*SodA-T_D-2_, and *Sa*SodM-T_D-2_, all of which exhibited increased catalysis with Fe, also showed elevated nOS_Fe_. The *Sa*SodA-L_D-2_-F_D-1_ variant exhibited significantly decreased nOS_Mn_, consistent with its reduced Mn-activity. The *Sa*SodM G_D-2_-L_D-1_ variant, which had increased Mn-preference, exhibited the opposite trend of increased nOS_Mn_ and significantly decreased nOS_Fe_.

After T_D-2_, the amino acid most frequently found at this position in Fe-preferring SodFM enzymes is valine^2^. We therefore introduced V_D-2_ mutations into each of the Mn-preferring enzymes, *Bs*SodA, *Lm*SodA and *Sp*SodA (CR=0.134, 0.296 and 0.384, respectively) by switching their X_D-2_/X_D-1_ residues for those of the Fe-dependent^34^ (Table 1) SodFM1 *Ba*SodA2 (V_D-2_-I_D-1_). In all cases, the variants exhibited increased Fe-preference (CR=1.487, 1.014 and 0.987, respectively; Table 1). These mutated *Bs*SodA, *Lm*SodA and *Sp*SodA variants also showed changes in spectra and nOS (Table 1), increasing nOS_Fe_ and decreasing nOS_Mn_, consistent with their cambialistic character.

X-ray crystallographic structural analysis of *Lm*SodA and *Ng*SodA demonstrated the anticipated architecture of their active sites (Supp. Fig. S2). Crucially, comparison of the structure of WT and the V_D-2_-I_D-1_ variant *Lm*SodA structures showed that their active site and hydrogen-bonding networks superimposed identically within the limits of crystallographic resolution (Fig. 3C), consistent with previous studies of the *Sa*SODs and their variants^1,2^. These data limit a role for substantial rearrangement of the active site in explaining the distinct metal preferences caused by these mutations. Instead, this is consistent with secondary coordination sphere residues tuning the redox properties of the metal ion through a mechanism not visible *via* X-ray crystallography with currently available resolution.

Finally, we constructed an X_D-2_/X_D-1_ mutant of the Fe-preferring SodFM2 isozyme, *Ng*SodB (T_D-2_-C_D-1_, CR=29.285), swapping them for the equivalent residues from the cambialistic^2^ SodFM2 from *Bacteroides fragilis* (*Bf*SodFM2 G_D-2_-F_D-1_, CR=0.728; Table 1). The *Ng*SodB G_D-2_-F_D-1_ was inactive and poorly expressed, in contrast with the active G_D-2_ single mutant, suggesting the amino acid at X_D-1_ can stabilise the presence of certain residues at the X_D-2_ position^1^. Despite a reduction in Fe-activity and CR, *Ng*SodB G_D-2_ remained a highly Fe-preferring enzyme (CR=7.783). Its nOS values also showed only minor, insignificant changes with respect to the wild type consistent with this slightly modulated metal-preference (Table 1).

Importantly, we observed that our variants either gained Fe activity and lost Mn activity, or gained Mn activity and lost Fe activity, indicating an inverse relationship between these two activities (Fig. 3D). The trend of simultaneous change in nOS and CR of these variants with mutations in the key X_D-2_ position that altered their metal preference (Fig. 3E) matched the pattern that was observed in comparisons of wild type variants (Fig. 2C). Together, these data are consistent with the hypothesis that metal-preference changes are dependent on modulation of metal-cofactor resting redox state, as measured by nOS.

### Role of classical neofunctionalization in the emergence of cambialistic SodM in *S. aureus*

We previously hypothesized that cambialistic *S. aureus* SodM had most likely emerged *via* neofunctionalization from its Mn-preferring SodA ancestor^1^, enabling *S. aureus* to resist Mn starvation during infection^8^. However, the exact evolutionary mechanism of this process remained unexplored. In our previous analysis of SodFMs sampled across the tree of life^1^, SodM homologues, found only in the *S. aureus/S. argenteus*-lineage, grouped at the base of SodA sequences from all oxidase-negative *Staphylococcus*, rather than with only SodA sequences from *S. aureus/S. argenteus*, as would be expected if they originated *via* gene duplication. Therefore, we repeated the analyses using 73 non-redundant (<98% pairwise nucleotide sequence identity) SodFM1s identified in all 21,452 available *Staphylococcaceae* genome assemblies. With this improved sampling, SodM and SodA from the *S. aureus/S. argenteus-*lineage grouped together in both nucleotide (Fig. 4A) and protein (Fig. 4B) phylogenies with high support (97% and 94% bootstrap support values, respectively), consistent with SodA and SodM emerging from ancestral gene duplication.

**Fig. 4.**
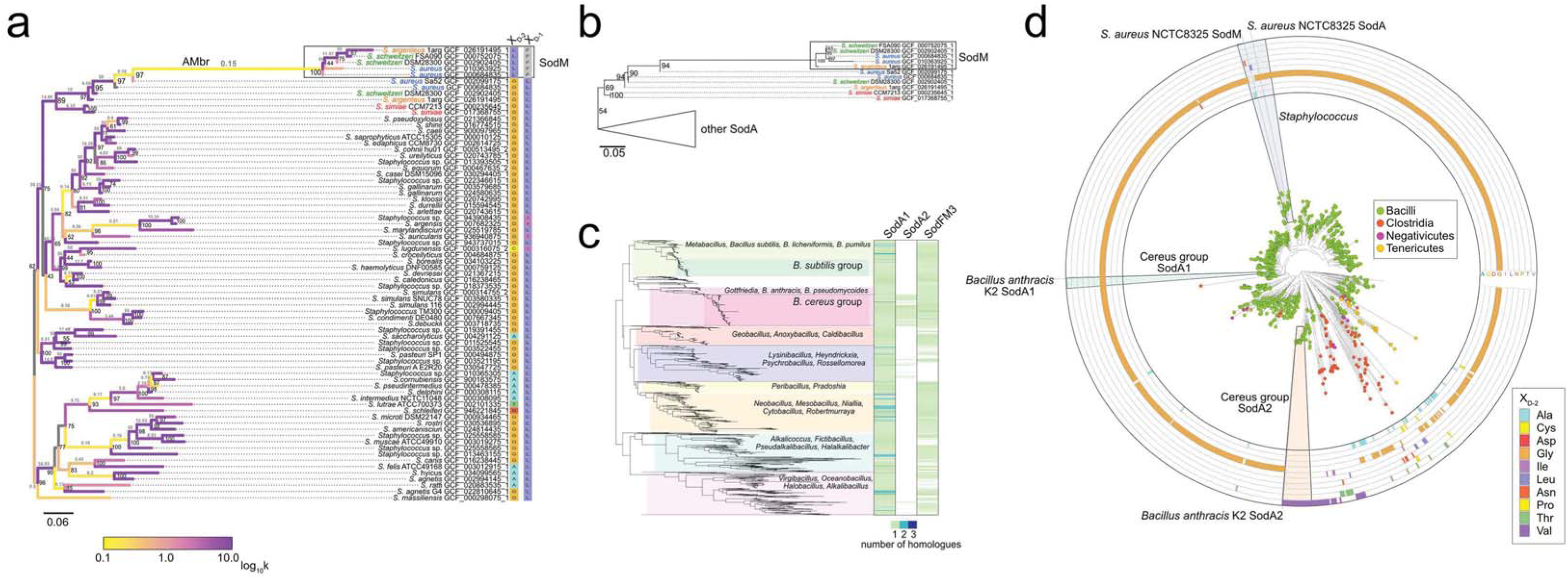
Investigating evolutionary mechanisms involved in SodFM1 metal-preference modulation. **A.** Maximum likelihood (GTR+F+R4 model) phylogeny of representative *sodFM1* nucleotide sequences sharing less than 98% identity sampled from all 21,452 available *Staphylococcus* genomes. All *sodM* sequences (black rectangle) from *S. aureus* (green), *S. argenteus* (orange), and *S. schweitzeri* grouped together with *sodA* sequences from the same species. Sequences of *sodA* from their closest relative (*S. simiae,* red) formed an outgroup, consistent with vertical inheritance of *sodA*. Branches of the tree were colour-coded with branch-dependent selection intensity parameter (k, grey values above branches). Values of k<1 (yellow) indicate relaxed selection and those of k>1 (purple) indicate intensified selection. **B.** Maximum likelihood (LG+G4) protein tree displaying topology consistent with that of the nucleotide tree (A). Support values (black, A and B) correspond to ultrafast bootstrap values from 1000 replicates. Scale bars represent the number of substitutions per site. C. Species tree of 1,115 non-redundant *Bacillaceae* genomes. Number of identified SodFM1 (SodA1 and SodA2) and SodFM3 homologues were mapped onto the tree (blue-green heatmap). D. Protein tree of SodFM1s sampled from across 828 representative Firmicutes and all available 11.394 *Bacillaceae* genomes. The identity of the X_D-2_ residues was mapped in the concentric circles. *S. aureus* SodM grouped together with SodA homologues, providing evidence for its emergence from a duplicated SodA ancestor. *B. anthracis* SodA2 grouped more closely with SodFM1s from Clostridia than *B. anthracis* SodA1 suggesting possible acquisition *via* LGT.

Next, we tested for evidence of the relaxation of selection following the inferred ancestral *sodA* duplication^35^. When SodM branches were tested against the background of all SodA, inferred relaxation (K=0.61) was significant (*p*=0.000, likelihood ratio=14.49). Mapping of the branch-dependent selection intensity parameter (k, Fig. 4A) onto the SodA/M phylogeny was consistent with *S. aureus/S. argenteus/S. schweitzeri* SodM and SodA being under intensified selection (k>1), while evidence of relaxed selection was mainly found on the long branch connecting SodAs and SodMs (AMbr, k=0.15). In an analysis excluding the AMbr, inferred selection intensification (K=2.08) was not significant (*p*=0.226, LR=1.46). This provided evidence in support of selection relaxation along the AMbr branch during the emergence of the last common SodM ancestor but not among the extant SodMs. Based on our biochemical and sequence analyses, position X_D-2_ clearly played a role in the evolution of cambialism in SodM^1,2^. Although they revealed evidence of episodic diversifying selection (*p*=0.000; BUSTED), neither of our analyses (*p*-value=0.67; MEME) provided any evidence for it acting on the residue X_D-2_. However, we did detect evidence for X_D-2_ being under pervasive negative selection (posterior probability [α>β] = 0.937; FUBAR) alongside 170 of 191 tested sites, consistent with a conserved role for both SodA and SodM in extant *S. aureus*. Previously we demonstrated that multiple different single mutations at this site can change biochemical properties of SodM towards higher Mn- or Fe-preference^2^. Therefore, negative selection acting at this site is consistent with our previous hypothesis that it was not simply higher Fe-preference but rather cambialism that was selected for in SodM^2^. Altogether, our results suggest that SodM evolved during a period of relaxed selection following the inferred duplication of the ancestral *sodA*, followed by intensified selection of its neofunctionalized descendent *sodM*, with likely retention of its cambialism through purifying selection.

According to the classical neofunctionalization model^36^, following a gene duplication, one copy remains under purifying selection while a second copy experiences a period of relaxed selection, which can facilitate acquisition of new properties that can be subsequently fixed. This classical model has been criticised and alternative models postulated^37–39^. However, the following empirical observations are consistent with SodM evolution having occurred according to the classical model: (1) although the ancestral capacity to function with Mn is diminished in cambialistic *Sa*SodM, it was never completely lost^1,2,8^; (2) SodM is capable of forming functional heterodimers with *Sa*SodA^8^; and (3) expression of *sodM* and *sodA* in *S. aureus* are differentially regulated^8^, which, if acquired early, could have enabled the two copies of the duplicated gene to experience different selection intensities. Taken together with our prior study^1^, this analysis establishes a likely evolutionary pathway for biological switching of catalytic metal-preference in a natural SodFM.

### Modulation of nOS underlies evolutionary changes of metal preference in natural SodFMs

Our data demonstrated that the SodFM resting spectral features and their resulting nOS values were predictive of the metal preference of natural SodFMs, and that secondary sphere mutations that altered SodFM metal-preference concomitantly altered nOS. The most recent identified evolutionary change in SodFM metal preference, the neofunctionalisation of the SodFM1 *Sa*SodM following duplication of the ancestor of Mn-preferring *Sa*SodA^1^, was associated with a change in nOS_Fe_ (Fig. 2). Next, we tested whether the modulation of nOS has played a conserved role in multiple other evolutionary metal-preference switches that we have inferred from our analyses of natural enzymes sampled across the tree of life^2^.

The only SodFM1s previously shown to be Fe-preferring are *B. anthracis*^34^ SodA2 and the SodFM1 from candidatus *Wolfebacteria*^2^ (c*W*SodFM1; CR=18.48) of the candidate phyla radiation (CPR). The CPR are a large and diverse phylum of bacteria that are poorly characterised and mostly uncultured, known primarily through their metagenome sequences. Like *Ba*SodA2, the c*W*SodFM1 enzyme also evolved from a Mn-preferring ancestor but through a different molecular pathway which also included change of the catalytic water-coordinating residue from Gln to His^2^. The c*W*SodFM1 also displayed nOS values consistent with its cofactor preference (Table 1). We constructed a G_D-2_ mutant of c*W*SodFM1, converting the valine residue in this position to a glycine, which is more typical of SodFM1 and generally indicative of Mn-preference^2^. This variant exhibited diminished Fe-activity without increased Mn-activity (c*W*SodFM1 V_D-2_G, CR=5.432) with a corresponding decrease in nOS_Fe_ (Table 1). Thus, isozymes from the predominantly Mn-preferring SodFM1 subfamily in which independent evolutionary alterations in metal-preference have been observed, creating either a cambialistic (*Sa*SodM) or an Fe-preferring enzyme (*Ba*SodA2 and c*W*SodFM1), exhibited consistent alterations in their spectral properties and nOS.

Next, we tested whether altered nOS was associated with changes in metal preference in other SodFM subfamilies. We characterised an atypical cambialistic^2^ (CR=0.728) *Bacteroides fragilis* isozyme (*Bf*SodFM2) from the predominantly Fe-preferring subfamily SodFM2. Both metal forms of *Bf*SodFM2 had high nOS values, consistent with its cambialistic activity (Table 1), analogous with cambialistic *Sa*SodM. We also created the T_D-2_-C_D-1_ variant of *Bf*SodFM2 in which we introduced residues from Fe-preferring SodFM2 *Ng*SodB. This mutation reverted *Bf*SodFM2 to its most likely ancestral Fe-preferring state, retaining high Fe-activity (CR=24.67) and nOS_Fe_ while strongly diminishing Mn-activity and nOS_Mn_ (Table 1).

Finally, we investigated a member of the SodFM3 subfamily. Isozymes from SodFM3 and SodFM4 have a more distant sequence, structural and phylogenetic relationship^2^ with the isozymes from SodFM1 and SodM2. Nonetheless, we observed nOS values for the Fe-preferring (CR=4.027) SodFM3 from *Akkermansia muciniphila* (*Am*SodFM3)^2^ that were consistent with those of the characterised Fe-preferring SodFM1 and SodFM2 enzymes (Table 1). This showed that the redox properties of these tetrameric SodFMs display characteristics similar to those observed in the dimeric enzymes of the SodFM1 and SodFM2 subfamilies.

The trend of nOS following metal-preference of activity was reproduced across the entire dataset (Fig. 3E). Collectively, these data demonstrate that all the examples of inferred evolutionary modulation of SOD metal preference identified so far^1,2^, including the ancient split between the SodFM subfamilies, and the more recent changes in metal-preference found in *S. aureus*, the bacilli, the *Bacteroides* and the CPR bacteria, all resulted in concomitant changes in nOS. These observations are consistent with the hypothesis that alterations in the redox properties of the active site apparent as changes in nOS play a conserved role in the evolutionary modulation of SOD metal preferences.

### Unexpected metal-binding by two SodFM1s with opposite metal-dependence

A pair of SodFM1s with different metal preferences in *B. anthracis* (*Ba*SodA1 and *Ba*SodA2)^34^ bears cursory resemblance to the pair identified in the staphylococci. We tested whether the SodA2 enzyme was acquired in an evolutionary process analogous to that described in *S. aureus*^1^. As previous data suggested that *Ba*SodA2 is highly Fe-preferring^34^, it is unlikely that it evolved in a process identical to that of *Sa*SodM as the complete loss of the ancestral Mn-activity could have posed challenges in maintaining the duplicated gene in the population. The scattered distribution of the SodA2 homologues found across the *Bacillaceae* species tree (Fig. 4C) is suggestive of multiple independent lateral gene transfer (LGT) events. The original source of the transferred genes is unclear but the protein tree of SodFMs sampled across Firmicutes (Fig. 4D) suggests that SodA2s may have originated from outside *Bacillaceae*. Grouping of the SodA2s with their closest *Clostridiaceae* homologues is nonetheless weakly supported (38% bootstrap value), reflecting their high level of sequence divergence. Therefore, the most parsimonious explanation is the acquisition of SodA2 *via* LGT in the ancestor of *B. anthracis/B. cereus*. However, the alternative hypothesis of emergence from the *Bacillus* SodA1 ancestor, followed by extreme sequence divergence that masks the phylogenetic signatures of SodA2’s true origin, cannot be fully excluded. Regardless of the mechanism, the result is a distinct SodFM1 repertoire in extant pathogenic species of the *Bacillus cereus*/*anthracis* group, including many pathogens, compared with their relatives including those from the lineage of non-pathogenic *B. subtilis* (Fig. 4C). Consistent with its high protein sequence homology to *Sa*SodA and *Bs*SodA (66% and 77% identical, respectively), *Ba*SodA1 shows canonical Mn-preference (Table 1) and likely reflects the state inherited from the last common *Bacillales* ancestor. However, while neofunctionalisation of *Sa*SodM created a cambialistic enzyme^1^, *Ba*SodA2 has undergone a more extreme shift in metal preference towards Fe (Table 1). Crucially, we observed metal-binding selectivity in *Ba*SodA2, as its Mn-loaded form could not be generated under our standard heterologous expression conditions or even using a range of *in vitro* unfolding/refolding dialysis protocols^34^ (Supp. Fig. S3). This precluded assessment of its nOS_Mn_ and its Mn-dependent catalytic activity and thus CR, but was consistent with a previous study that demonstrated *Ba*SodA2 was associated exclusively with Fe in its native cytosol^34^. Importantly, while *Ba*SodA1 exhibited typical spectra and nOS values expected from a Mn-preferring SodFM1, *Ba*SodA2 exhibited nOS_Fe_=0.35, consistent with its characterisation as an Fe-active enzyme (Supp. Fig. S1; Table 1). These atypical properties reflect the high level of SodA2 sequence divergence and can potentially explain the difficulty in inferring its origins. Furthermore, they may also explain the relatively high frequency of sharing SodA2s *via* LGT in *Bacillaceae*, as opposed to, for example, acquisition of more common Fe-preferring SodFM2s or SodFM3s. It is possible that selectivity for Fe binding may be more difficult to achieve in SodFMs, and that it can provide some unique selective advantages. These data also pose a question about the evolutionary mechanism that could lead to emergence of an Fe-selective enzyme from a highly Mn-preferring ancestor, and why such selectivity is not more widely observed among strongly metal-preferring SodFMs. Neofunctionalisation cannot be completely excluded as we previously observed a complete switch between Fe-preference and Mn-preference *via* a single mutation in *Wosearchaeota* SodFM4 (V_D-2_ to A_D-2_ mutation)^2^. However, no such mutations are known to be possible in Mn-SodFM1s thus far. Therefore, given our current knowledge, subfunctionalisation seems a more likely mechanism, suggesting the existence of at least one intermediate cambialistic ancestor of SodA2 prior to the emergence of Fe-specificity and/or Fe-selectivity.

Reciprocal double mutagenesis at X_D-2_/X_D-1_ of the *Ba*SodFM1s was performed (G_D-2_-L_D-1_ in Mn-preferring *Ba*SodA1, CR=0.196; V_D-2_-I_D-1_ in Fe-active *Ba*SodA2, CR=unmeasurable) to investigate if these residues can play a role in switching metal preference in the bacilli, as in the staphylococci. The shift in metal preference towards cambialism in the *Ba*SodA1 V_D-2_-I_D-1_ mutant (CR=0.743) was analogous to that observed in mutagenesis of *Sa*SodA, as were its significant changes in nOS (Table 1). The reciprocal mutation, inserting the G_D-2_-L_D-1_ residues of *Ba*SodA1, which are typical of SodFM1s and generally indicate Mn-preference^2^, into *Ba*SodA2 created an unstable and inactive variant. A G_D-2_ single mutant remained Fe-active when analysed by in-gel assay from cell lysates, however neither variant could be purified in sufficient quantity to analyse their spectra. Therefore, we conclude that the challenging adaptation to Fe-activity in this SodFM1 has made its active site less able to switch between metal-preferences through this mutagenesis strategy. This is consistent with our previous data for c*W*SodFM where a second mutation of water-coordinating His to Gln was required to increase its Mn-dependent activity, and for the SodFM2 from *Agrobacterium tumefaciens* that we were unable to switch to its most likely ancestral Mn-preference^2^. This suggests the presence of secondary epistatic mutations in SodFMs subjected to the most extreme evolutionary changes is required to induce metal-preference modulation (*At*SodFM2, *cW*SodFM1 *Ba*SodA2), and to cause metal selectivity (*Ba*SodA2) while retaining specific activity and protein fold stability.

Notably, *Ba*SodA2 was not the only isozyme that displayed unanticipated metal binding during recombinant production. Despite its widespread adoption as a ‘model’ Mn-preferring SodFM1 for decades^18,23,40^, we observed here, as previously^2^, that *Ec*SodA has unusual properties. Under our standardised expression conditions in minimal medium, essential to control metal-loading of SodFM isozymes, *Ec*SodA was the only SodFM tested that did not bind Fe inside *E. coli* cells cultured aerobically in the absence of Mn. Under these conditions, *Ec*SodA was expressed only at low levels and resulting protein preparations lacked significant Fe^2^. This observation is particularly striking as we were able to produce multiple SodFMs from all subfamilies sampled from across the tree of life in the same system^2^, but it was the homologously expressed *E. coli* SodA that displayed unusual properties, suggesting that it may represent a specific adaptation. Synthesis of the Fe form of *Ec*SodA required the use of a rich growth medium supplemented with additional Fe. Under these conditions, the metal-loading of the isozyme could not be fully controlled and was loaded with mixtures of Fe and Mn, which only allowed a limited spectral analysis to be performed. Crucially, a preparation of *Ec*SodA that contained mostly (∼80%) Fe exhibited a spectrum consistent with the other characterised Mn-preferring SodFM1s (Supp. Fig. S1).

We reciprocally interconverted the two second sphere residues in the *E. coli* SodFM1/SodFM2 pair (G_D-2_-L_D-1_ in Mn-preferring *Ec*SodA, CR=unmeasurable; T_D-2_-V_D-1_ in Fe-preferring *Ec*SodB, CR=27.51). *Ec*SodB G_D-2_-L_D-1_ showed reduced Fe-activity and a CR shift towards cambialism but remained Fe-preferring (CR=3.992), and consequently exhibited only small changes in nOS that were not significant (Table 1). *Ec*SodA T_D-2_-V_D-1_ was still a highly Mn-active enzyme (∼70% of wild type activity) with a high nOS_Mn_ (Table 1). However, like the wild type, the Fe-loaded form of *Ec*SodA T_D-2_-V_D-1_ could not be generated under our standard conditions in sufficient yield for spectroscopic analysis, suggesting different sites are responsible for its unusual metal-binding properties. Taken together, the *E. coli* SodFMs, unlike many of their SodFM1 (*Ec*SodA) and SodFM2 (*Ec*SodB) relatives, display a high degree of resistance to metal-preference alteration *via* the common metal-preference modulation pathways involving mutation of the X_D-2_-X_D-1_ residues^2^.

In summary, two SodFMs from *E. coli* and *B. anthracis*, uniquely among all the isozymes we have biochemically tested, exhibited a degree of metal selectivity under the tested conditions. Whereas all other SodFMs are competent to bind both metal cofactors, regardless of which confers catalysis, these enzymes displayed binding to only one of the respective metals, Mn or Fe, when expressed inside aerobically grown Δ*sodA*Δ*sodB E. coli* cells under defined metal culture conditions. Notably, SodA is already present within the manganese-poor cytosol of *E. coli* prior to an oxidative stress response^41^, and emergence of a level of metal-selectivity might prevent its mis-metalation. If so, a need for *B. anthracis* to have *Ba*SodA2 already present in its manganese-rich cytosol^42^ could also be hypothesised. Remarkably, two distinct isozymes with a degree of metal-selectivity have evolved within the SodFM1 subfamily from most likely Mn-preferring, metal-nonselective ancestors, one with retention of the ancestral metal preference (Mn-selective *Ec*SodA) and the other with a metal-preference switch (Fe-selective *Ba*SodA2). These enzymes could prove to be useful models to study the mechanisms of metal binding in SodFMs.

### The nOS reflects the enzymatic activity and resting redox poise of SodFM cofactors

Comparison of the nOS values derived from spectral analyses of ten wild type isozymes and nine mutated variants illustrated an intriguing trend (Fig. 3E). Metalated forms of all SodFMs that exhibited significant catalytic activity also possessed substantial nOS values (generally 0.25<nOS<0.95). This indicates that all of these catalytically active samples contained a significant population of oxidised metal cofactors at rest. Conversely, all forms of SodFMs that exhibited negligible catalytic activity possessed nOS values close to zero, consistent with their metal ions being completely reduced at rest.

To determine the relationship between metal preference and nOS for each metal cofactor, we plotted each set of nOS values, nOS_Fe_ and nOS_Mn_, from our replicated spectral analyses of SODs against their CR (plotted on a logarithmic scale; Fig. 5). This clearly illustrated that metal preference is a continuum between perfect metal specificity extremes (here represented by the more metal-selective *Ec*SodA and *Ba*SodA2, whose nOS values could not be fully assessed), rather than the traditional view of three discrete states (MnSOD, FeSOD, camSOD). Trendline fitting of these datasets yielded a striking observation; nOS_Fe_ is proportional, while nOS_Mn_ is inversely proportional to CR (Fig. 5). The resulting trendlines intersected at CR=0.97 and nOS=0.27, representing the anticipated properties of a hypothetical, near-perfectly cambialistic isozyme. Furthermore, this trend still broadly held when additional data, acquired from the more highly divergent SodFMs and their mutants (Fig. 3E), were included (Supp. Fig. S4).

**Fig. 5.**
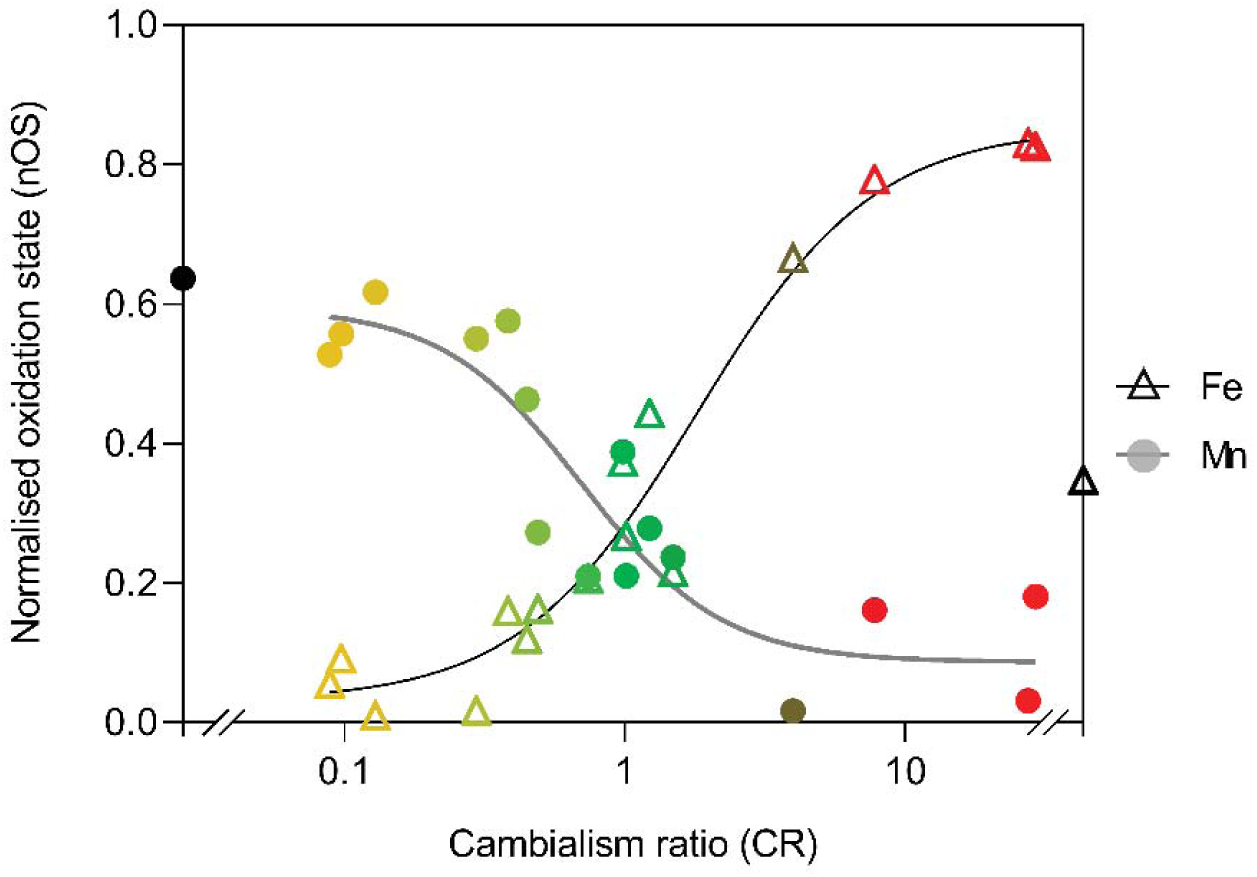
Inverse trends of cambialism ratio and oxidation state in Mn-loaded and Fe-loaded SodFMs. When nOS values for Fe-loaded (open triangles) and Mn-loaded (closed circles) forms of all the SodFM isozymes analysed in triplicate were plotted against CR on a logarithmic scale, we detected two inverse trends. The enzymes with low activity have very low oxidation at rest, while those with increased activity displayed higher oxidation at rest, for both metal-forms. Data points are coloured by cambialism ratio using a tricolour gradient from yellow (Mn-preferring) to green (cambialistic) to red (Fe-preferring). Trendlines were calculated in GraphPad Prism (Version 9.3.1) using a sigmoidal 4-parameter logistic model. The R^2^ values of the non-linear fit to the data were R^2^ = 0.957 for the Fe-loaded enzymes and R^2^ = 0.841 for the Mn-loaded enzymes. As a real CR could not be calculated for *Ba*SodA2 due to apparent metal-discriminating selectivity of binding, this has been artificially assigned to the extreme of the CR scale, with its single nOS value represented by a black symbol, which was not used in the calculation of trendlines.

Taken together, the trend observed between cambialism ratio and nOS of all tested isozymes is strongly indicative that they are intrinsically linked features of SodFMs. The diverse SodFMs analysed here represent a spectrum of evolutionarily divergent Fe/Mn SODs and include examples shown to have undergone ancient or more recent evolutionary modulations in preference^2^. This argues that this relationship between the metal-preference and the cofactor’s resting redox properties underlies the evolutionary mechanism of metal-preference switching. We propose that this relationship is determined by the effects of the protein architecture on the biophysical properties of the metal cofactor within this metalloenzyme family, mediated by the ion’s secondary coordination sphere, and that evolution must alter nOS in order to modulate metal-preference.

## Discussion

Metalloproteins are ubiquitous in biology, with metals having essential catalytic or structural functions in enzymes from all six Enzyme Commission classes. Metalloenzymes comprise nearly half of all oxidoreductases, in which *d*-block metals confer redox activity^6^. It is generally assumed that such redox metalloenzymes are highly specific for their cognate metal^11–13^ because they optimise their active site to manipulate the metal’s reduction potential to achieve a suitable midpoint potential for efficient catalytic cycling. This optimisation of potential by the protein architecture is called redox tuning. Due to inherent differences in electronic configurations of different metals, the tuning imparted by a protein structure on its cognate metal ion (e.g. Fe) will be inappropriate for tuning the distinct inherent potential of a non-cognate metal cofactor (e.g. Mn). Such a model would explain why there is only a limited number of characterised examples of natural metalloenzymes that exhibit cofactor-flexibility (cambialism), although recent analyses suggest they may be more common in nature than previously thought^2,8,13,43–45^.

For a metalloenzyme that is optimised for one redox metal cofactor to evolve an alternative metal specificity, under suitable selection pressure, this model requires re-optimisation of its active site for the new metal. This process could enable the new metalloenzyme to gain an entirely new catalytic function using the new cofactor, an idea supported by numerous bioinformatic observations in diverse protein fold families^3,5,46^. Alternatively, it could result in a switch to a new metal specificity to achieve the same chemical reaction, of which there are far fewer examples^43,44^ but is a rational adaptation to changes in environmental metal availability^1^. The molecular mechanisms by which tuning is achieved by the protein architecture is unclear, however, and thus we lack an understanding of how tuning can be re-shaped by evolution to change enzymatic metal-preference. Furthermore, it is unknown whether such changes in metal utilisation must pass through intermediate evolutionary stages that have sub-optimal tuning, resulting in reduced efficiency but potentially with some level of cambialism with both the ‘old’ and ‘new’ cofactors. SodFMs are a useful model system with which to interrogate these important evolutionary questions.

We exploited the absorption properties of SodFM cofactors, the intensities of which report on the oxidation state of the bound metal ions, to assess whether the redox properties of SodFMs followed their trend in catalytic metal-preference. By sampling diverse SodFMs from across the CR range, we demonstrated an inverse relationship between nOS_Fe_ and nOS_Mn_ in natural enzymes. In all cases, enzymatically active metal-loaded forms exhibited nOS values consistent with their population of molecules containing a mixture of both oxidised and reduced metal cofactors at aerobic equilibrium. Conversely, the forms lacking significant activity exhibited nOS values close to zero, implying they contained exclusively reduced metal cofactors. On introducing mutations into the cofactor’s secondary coordination sphere that modulated SodFM metal-preference, we observed concomitant changes in spectra and nOS consistent with the hypothesis that the mutation-induced metal-preference changes were dependent on modulation of the metal cofactor’s resting redox state. These data demonstrate unequivocally that both the nOS and the metal-preference of a SodFM are manipulated through changes to the cofactor’s secondary coordination sphere. Finally, we tested whether the same nOS modulation was observed during natural evolution of metal preference by characterising some of the most divergent SodFMs known: those that have evolved extreme metal-preference within the context of their close relatives (e.g. c*W*SodFM1, *Bf*SodFM2)^2^, a member of one of the most highly divergent subfamilies (*Am*SodFM3)^2^, and two isozymes that exhibit a degree of metal-binding selectivity (*Ec*SodA and *Ba*SodA2). All SodFM isozymes and variants tested followed the same trend in nOS relative to CR. Taken together, the data presented here suggests that SodFM metal preference naturally evolves through a mechanism that is inextricably linked to changes in the metal’s oxidation state at rest, mediated by the cofactor’s secondary coordination sphere.

The redox tuning model was proposed to explain the differing cofactor specificity between the pair of isozymes from *E. coli* and has some empirical support in that system^21–23^. Notably, however, *Ec*SodA was the only isozyme tested herein that was not metalated with Fe when expressed homologously inside aerobic *E. coli* under our standard expression conditions in minimal medium, indicating it displays unusual strong Mn-selective metal binding properties within this system. This unusual behaviour of *Ec*SodA should give caution in interpreting the body of biochemical studies on this enzyme^18,21,40,47^, as they may not be generalisable to the whole SodFM family. Interestingly, selectivity towards the other metal cofactor, Fe, was observed in another SodFM1, *Ba*SodA2, which was unable to bind Mn *in vitro* or *in vivo*. These two isozymes are the only SodFMs yet studied that display such metal-binding properties, a phenomenon which warrants further mechanistic investigation. These observations also raise a question why such metal selectivity is not more widespread, at least among highly metal-preferring SodFMs where mis-metalation leads to inactive isoforms. Both of these enzymes can serve as models for future studies of the biochemical mechanisms and physiological selection pressures enabling and driving the novel emergence of metal-selectivity in SodFMs.

The overall trends between the resting spectra and the associated nOS of a SodFM and its catalytic metal preference were robust, assembled from biochemical and biophysical measurements obtained from more than 100 separate preparations of purified recombinant protein, and are consistent with prior observations in the literature^31,48,49^. The inverse relationship between nOS values suggests that evolutionary changes in metal-preference of SodFMs follow a predictable biochemical path. For a highly Mn-preferring isozyme to gain increased catalysis with Fe, it must increase nOS_Fe_. However, because these parameters are intimately linked, increasing nO_Fe_ will necessarily result in decreased nOS_Mn_, thereby reducing its activity with Mn. These properties can be balanced within cambialistic enzymes, in which nOS_Mn_≈nOS_Fe_ and CR≈1. This process occurred in the evolution of *Sa*SodM during its divergence from a Mn-preferring ancestor^1^, likely under selection pressure of decreased Mn availability within the mammalian host^8^. The converse evolutionary process, increasing nOS_Mn_ at the expense of nOS_Fe_, likely yielded the cambialistic *Bf*SodFM2 from its highly Fe-preferring ancestor^2^. Larger changes in nOS can more dramatically switch metal-preference, resulting in larger changes in CR, such as those that occurred in the unique Fe-preferring SodFM1 from candidatus *Wolfebacteria* or the Mn-preferring SodFM2 from *Agrobacterium tumefaciens*^2^. It appears that even more extreme changes must have occurred within *Ec*SodA, and especially *Ba*SodA2, resulting in altered metal-binding selectivity. It remains to be studied whether these changes in metal-preference in *Bf*SodFM2, *Ec*SodA or *Ba*SodA2 are specific evolutionary adaptations, and if so, under what selection pressure they emerged. Although it is not necessary for a full evolutionary switch in preference, i.e. from a Mn-preferring SodFM to become Fe-preferring or vice versa, to transition via a cambialistic evolutionary intermediate, this is likely in at least some cases. Future studies should seek to verify whether cambialism itself has been selected, as suggested in *S. aureus*^2^, rather than being a transitional evolutionary state, and to determine whether the changes in redox properties observed here represent a conserved mechanism by which evolution of catalytic metal-preferences occur in other redox metalloenzymes.

## Materials & Methods

### Bacterial strains and culture conditions

The Fe/Mn SOD-deficient *E. coli* strain, BL21 (λDE3) Δ*sodA*Δ*sodB*, was used as expression host to eliminate *Ec*SodA/*Ec*SodB contamination from recombinant protein preparations^2^. *E. coli* strain DH5α was used for molecular biology. BL21 pLysS Δ*sodA*Δ*sodB* was used to improve expression of the wild type and mutant variants of *Ba*SodA2, which expressed poorly in BL21 Δ*sodA*Δ*sodB*. For selection, 100 μg mL^−1^ ampicillin or 50 μg mL^−1^ kanamycin were added to the growth media.

### Molecular biology

Cloning of the pET22b constructs for expression of *Ba*SodA1, *Ba*SodA2, *Lm*SodA, *Sp*SodA, *Ng*SodB, *Ec*SodA (from *E. coli* B), *Ec*SodB, c*W*Sod, *Am*Sod, and *Bf*Sod was previously described^2^. Constructs for the heterologous expression of the genes encoding *Bs*SodA and the wild type and mutant variants of *Sa*SodA and *Sa*SodM in pET29a produced in a prior study^1^ were sub-cloned into pET22b(+), except for those used for expression of the *Sa*SodA T_D-2_ and *Sa*SodM T_D-2_ mutated constructs, which remained in pET29a. The genes for sub-cloning were excised from their vector through *Nde*I/*Sac*I (NEB) restriction, purified after electrophoresis (1% w/v agarose) using a gel extraction kit (Sigma-Aldrich), and ligated (T4 Ligase, NEB) into *Nde*I/*Sac*I-digested, Antarctic phosphatase-treated (NEB) pET22b(+) vector. *E. coli* DH5α transformants were ampicillin-selected and screened by PCR using T7 primers. Sequences of the inserts of all constructs were confirmed through DNA sequencing (Eurofins).

### Mutagenesis

Primers for site-directed mutagenesis were designed using the NEBaseChanger tool (NEB) and synthesised (Sigma and IDT). Mutagenesis was performed by PCR using the appropriate primer pairs, treated with KLD enzyme mix (NEB), then used to transform competent *E. coli* DH5α cells. Transformants were screened by colony PCR. Sequences were confirmed through DNA sequencing.

### Expression and purification of recombinant SODs

Plasmid constructs were transformed into chemically competent *E. coli* BL21(λDE3) Δ*sodA*Δ*sodB* cells and selected on LB agar plates containing ampicillin (pET22) or kanamycin (pET29). A single transformant was inoculated into 50 mL selective media (M9 or LB) and cultured overnight at 37°C with 180 rpm orbital shaking. The pre-culture was then used to inoculate 0.5–1.0 L of fresh selective M9 media supplemented with 1% (w/v) glucose to an OD_600nm_ ∼0.05. For production of Fe-enriched samples of *Ec*SodA only, the cells were cultured in LB medium supplemented with 100 μM Fe. Protein expression was induced by addition of 1 mM isopropyl-β-D-1-thiogalactopyranoside (IPTG), accompanied by addition of either 300 μM ammonium iron sulfate or 1 mM MnCl_2_, followed by incubation at 37°C for 4 h with 180 rpm orbital shaking. Cells were harvested by centrifugation (4,200 *g*, 25 min, 4°C), washed in 20 mM Tris pH 7.5 and frozen at -20°C.

Cell were lysed by sonication on ice in 20 mM Tris pH 7.5, 1x cOmplete EDTA-free protease inhibitor (Roche), 100 μg mL^−1^ lysozyme, 10 μg mL^−1^ DNase, followed by centrifugation at 19,000 g, 4°C. Cleared cell lysate was subjected to a two-step chromatographic separation using an AKTA (GE Healthcare). Recombinant SODs were initially purified using anion exchange chromatography (Hi Trap Q HP column, GE Healthcare) in 20 mM Tris pH 7.5 buffer with 0–1 M NaCl gradient elution. Eluent fractions containing SODs were identified by SDS-PAGE, and were pooled and concentrated to a final volume of 1 mL using centrifugal filtration devices (Amicon, 10 kDa cutoff) for subsequent size exclusion chromatography (SEC). For *Ec*SodA, an additional cation exchange step (Hi Trap SP FF column, GE Healthcare) was performed in 20 mM MES pH 5.5 buffer with 0–1 M NaCl gradient. *Ec*SodA eluted in the flow-through of both ion exchange steps and was dialysed overnight at 4°C in 20 mM Tris pH 7.5, 150 mM NaCl before concentrating for size exclusion chromatography (SEC). SEC was performed in 20 mM Tris pH 7.5, 150 mM NaCl buffer on a Superdex 200 16/600 GL column (GE Healthcare). SDS-PAGE confirmed that SODs eluted in a single peak at 79-86 ml. Protein concentration of recombinant protein preparations was determined by absorbance at 280 nm using each enzyme’s theoretical extinction coefficient (ε, ProtParam), and metal loading was analysed by inductively coupled plasma mass spectrometry (ICP-MS) or inductively coupled plasma optical emission spectrometry (ICP-OES). Protein preparations were split into 0.2–1.0 mL aliquots and stored at -20°C.

Where necessary, for poorly Mn-loaded protein preparations, metal exchange was achieved by unfolding SOD proteins and refolding through dialysis in the presence of 10 mM MnCl_2_ as previously described^1^. After dialysis, protein was concentrated using centrifugal filtration devices to 0.5 mL, and purified by SEC (Superdex 200 Increase 10/300 column, GE Healthcare) to confirm oligomeric state and remove any aggregated protein and unbound metal ions.

### SOD activity assays

SOD activity was assessed qualitatively using a gel-based negative staining assay^8^, or quantitatively using an adaptation of this method in a 96-well plate format, as previously described^2^.

### UV/visible spectroscopy

Absorption spectroscopy was performed on a Perkin-Elmer λ35 spectrophotometer, collecting data at room temperature between 200 and 700 nm using quartz cuvettes with a path length of 10 mm (ALS). Measurements of resting spectra (representing the oxidation status of the purified protein when equilibrated under aerobic conditions) were performed in 20 mM Tris pH 7.5, 150 mM NaCl, with samples prepared at 100 μM protein. Each resting sample was split into two, and each half was either chemically oxidized or reduced by incubation with 1 molar equivalent of potassium permanganate or 3 molar equivalents sodium dithionite for 10 min, respectively, followed by extensive buffer exchange using centrifugal filtration in 20 mM Tris pH 7.5, 150 mM NaCl. Spectra were converted to extinction coefficient, ε, using protein concentration and were normalised to nOS for comparison between isozymes as follows:

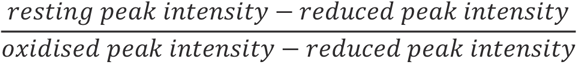

where peak intensity corresponds to absorbance at 480 nm or 350 nm for Mn-loaded and Fe-loaded enzymes, respectively. In order to test whether the resting state could be enzymatically regenerated after complete oxidation or reduction, the buffer-exchanged oxidised and reduced forms of SOD were mixed in an equimolar ratio and incubated with 0.1 mM riboflavin for 20 min on a white light box, before extensive buffer exchange to remove the riboflavin. UV-visible spectra showed that the resting state was regenerated, demonstrating enzymatic activity had not been lost through chemical oxidation or reduction.

### Elemental analysis

Aliquots of purified protein samples (20 μM) in 20 mM Tris pH 7.5, 150 mM NaCl were each diluted to 5 ml with 2% HNO_3_ for elemental analysis. Elemental composition of the resulting acid solutions was quantified using an iCAP RQ ICP-MS (Thermo Fisher Scientific) instrument (University of Plymouth Enterprise Ltd), or using an iCAP Pro ICP-OES (Thermo Fisher Scientific) instrument, as previously described^2^.

### X- ray crystallography

Protein preparations for crystallography were concentrated to 15-19 mg mL^−1^ and were subjected to crystallisation screening using a Mosquito liquid handling robot (TTP Labtech) with commercially available matrix screens: PACT, JCSG+, Structure, Morpheus (Molecular Dimensions) and Index (Hampton Research) in 96-well MRC crystallization plates (Molecular Dimensions) using the sitting drop of vapour-diffusion method (two drops per well, containing 100 nL + 100 nL and 200 nL + 100 nL of protein and crystallisation liquor, respectively), incubated at 20 °C. *Ng*SodB crystallised in Morpheus condition D1 (0.1 M MES/imidazole pH 6.5, 10% w/v PEG 20000, 20% v/v PEG MME 550, 0.02 M of each additive alcohol: 1,6-hexanediol, 1-butanol, (RS)-1,2-propanediol, 2-propanol, 1,4-butanediol, 1,3-propanediol), *Lm*SodA crystallised in PACT condition A8 (0.2 M Ammonium chloride, 0.1 M Sodium acetate pH 5.0, 20% w/v PEG 6000) and *Lm*SodA V_D-2_-I_D-1_ crystallised in Structure condition D6 (0.2 M Ammonium sulfate, 30 % w/v PEG 8000). Crystals were harvested in 20% PEG-400 cryoprotectant and flash-frozen in liquid nitrogen. X-ray diffraction data were collected at the Diamond Light Source synchrotron (Didcot, UK) on beamline I03 and beamline I24 at 100 K on two trips: mx24948-132 and mx24948-124. Structural solution and model building were as described previously^2^. 3LIO was used as a search model for molecular replacement for *Ng*SodB and 2RCV was used as a search model for *Lm*SodA. The generated structural model of *Lm*SodA was subsequently used to solve the structure of the *Lm*SodA mutant variant.

### Evolutionary analysis of Staphylococcus SodFM1 homologues

All 21,452 available NCBI RefSeq *Staphylococcaceae* genome assemblies (excluding atypical) were downloaded on 12.01.2024. SodFM homologues were identified using PFAM^50^ hidden Markov model (HMM) profiles (Sod_Fe_C, PF02777.21; Sod_Fe_N, PF00081.2) with hmmsearch^51^ (-E 1e^−5^) profile search implemented in HMMER3.3. HMM search identified 37,603 *Staphylococcus* SodFM homologues including 478 unique protein sequences encoded by 1051 unique nucleotide sequences. Of the 1051 nucleotide sequences, 73 shared less than 98% pairwise sequence identity (mean of 86% for protein and 83% for nucleotide sequences) with each other and only these 73 sequences were used in the subsequent analyses. The proteins sequences were aligned with MAFFT^52^ and corresponding nucleotide alignments were generated with local version^53^ of PAL2NAL_v14.

Pairwise sequence identity was calculated with trimAl^54^ (-sident). Phylogenies were reconstructed using IQ-TREE^55^ with 1,000 ultrafast bootstraps^56^. Nucleotide *Staphylococcaceae* SodFM tree was generated under best fitting GTR+F+R4 model, and protein *Staphylococcaceae* SodFM tree was generated under best fitting LG+G4 model selected with ModelFinder^57^ according to Bayesian information criterion implemented in IQ-TREE. Bootstrap values and branch lengths were removed in R using ape library^58^ and the phylogenies and the corresponding nucleotide alignments were used in the subsequent analyses. Positively and negatively selected sites were inferred with FUBAR^59^. Gene-wide positive selection at least one site on at least one branch was tested with BUSTED^60^. Specific sites under episodic positive selection were inferred with MEME^61^.

RELAX^35^ analysis was used to test trends in the relaxation or intensification of the strength of natural selection along specified set of test branches relative to the specified background branches.

### Bioinformatic identification of Bacillus SodFM homologues

All 11.394 available NCBI RefSeq Bacillaceae genome assemblies (excluding atypical) were downloaded on 12.12.2023. HMM searches with hmmsearch (-E 1e^−5^) identified 23,278 SodFM homologues in *Bacillaceae* including 8,958 SodFM3 and 14,320 SodFM1 redundant sequences (i.e. including identical sequences encoded across e.g. multiple *B. anthracis* genome assemblies). The identified SodFMs included 2,740 unique protein sequences of which 1,137 were SodFM3s and 1,603 SodFM1s. SodFM1 subfamily was subdivided into 1,250 SodA1 (9,868 redundant) and 353 SodA2 (4,452 redundant) unique protein sequences based on their protein tree topology and Xd-2 residue identity verified in MAFFT alignments.

*Bacillaceae* species tree was generated using concatenated alignment of seven universally conserved orthologues as described previously^2^. To reduce the oversampling of highly identical genome assemblies from some lineages (e.g. multiple *B. cereus/anthracis* isolates), only those containing unique sequences of all seven universally conserved orthologues were used, resulting in the final dataset of 1,115 genomes encoding 111 SodA2s, 1,185 SodA1s, and 819 SodFM3s.

Firmicutes dataset included 828 representative genome assemblies sampled from across all major Firmicutes lineages. HMM searches with hmmsearch (-E 1e^−5^) identified 2,153 (2,759 total) unique SodFMs including 1,359 unique (1,772 total) SodFM1s in Firmicutes genomes. *Bacillaceae* and Firmicutes SodFM protein trees were generate in using IQ-TREE with 1,000 ultrafast bootstraps under WAG+G4 model.

### Other bioinformatics methods

Multiple sequence alignments were inspected using Jalview^62^. Phylogenetic trees were inspected in Archaeopteryx^63^. The phylogenies were visualised and annotated in R with ggplot2^64^, ggtree^65^, ape^58^, treeio^66^, ggtreeExtra^67^, and ggnewscale.

---

**Supplementary Figure S1.**
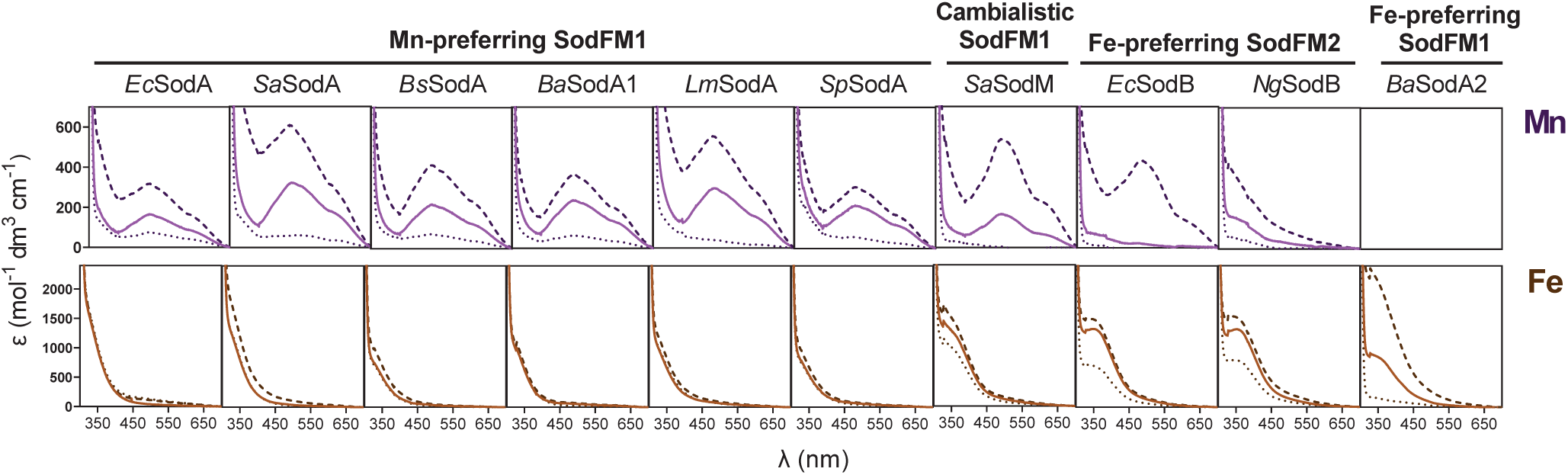
Spectra of wild type SodFMs at rest. UV-visible absorption spectra of all wild type SodFMs analysed in triplicate in this study when Mn-loaded (upper panels, purple lines) and Fe-loaded (lower panels, brown lines). Resting spectra of proteins equilibrated at atmospheric conditions are shown as a solid line, spectra of oxidised and reduced proteins are shown as darker dashed and dotted lines, respectively. This analysis demonstrated that the resting spectra lie between oxidised and reduced spectra for catalytically active forms (i.e. when Mn-bound for Mn-preferring SodFM1s, when Fe-bound for Fe-preferring SodFM2s, and in both metal forms for cambialistic *Sa*SodM) but are similar to the reduced spectra for inactive forms. Spectra are shown as *n* = 1, but are representative of further replicates (*n* = 3). As we were unable to generate the Fe-loaded form of *Ec*SodA nor the Mn-loaded form of *Ba*SodA2, no corresponding spectra were obtained for these forms.

**Supplementary Figure S2.**
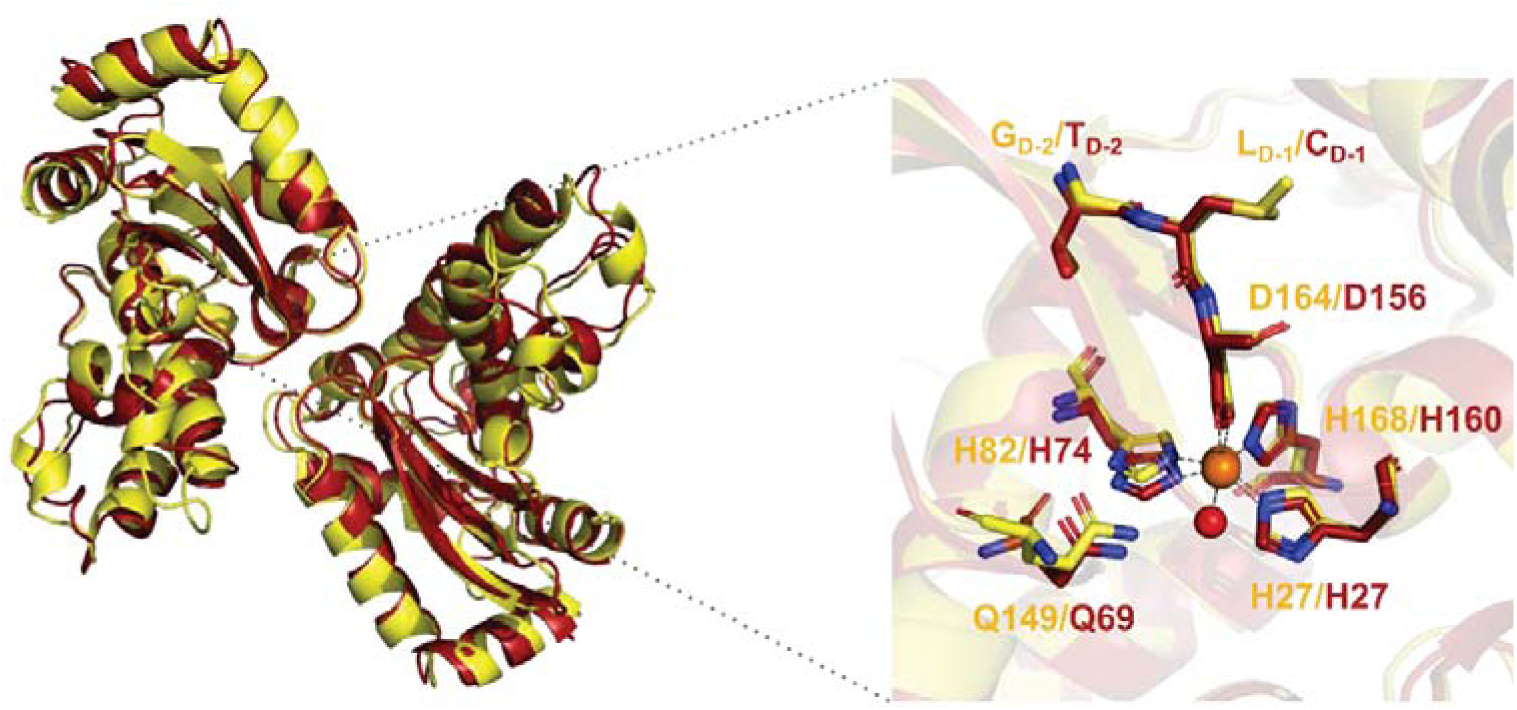
Structural comparison of the SodFM1 subfamily isozyme, *Lm*SodA, and the SodFM2 subfamily isozyme, *Ng*SodB. Overlaid structural models derived from X-ray crystallography showing the homodimeric assemblies (left) and zoomed in view of the active sites (right panel) of the Mn-preferring SodFM1 isozyme from *Listeria monocytogenes* (yellow) and the Fe-preferring SodFM from *Neisseria gonorrhoeae* (red). Within the active sites (right), key amino acid residues are shown in ball-and-stick representation: the metal binding ligands (H27 in both, H82 in *Lm*SodA and H74 in *Ng*SodB, H168 in *Lm*SodA and H160 in *Ng*SodB, and D164 in *Lm*SodA and D156 in *Ng*SodB), the water-coordinating residue (Q149 in *Lm*SodA and Q69 in *Ng*SodB), and the two second sphere residues that were targeted for mutagenesis herein, X_D-2_ (G162 in *Lm*SodA and T154 in *Ng*SodB) and X_D-1_ (L163 in *Lm*SodA and C155 in *Ng*SodB). The metal ions are shown as orange spheres and a solvent molecule is indicated with a red sphere. Despite these isozymes being from distinct subfamilies (47.9% sequence identity), their overall structures are remarkably similar, including the position of an important Gln residue which is encoded in distinct sequence loci in these two subfamilies.

**Supplementary Figure S3.**
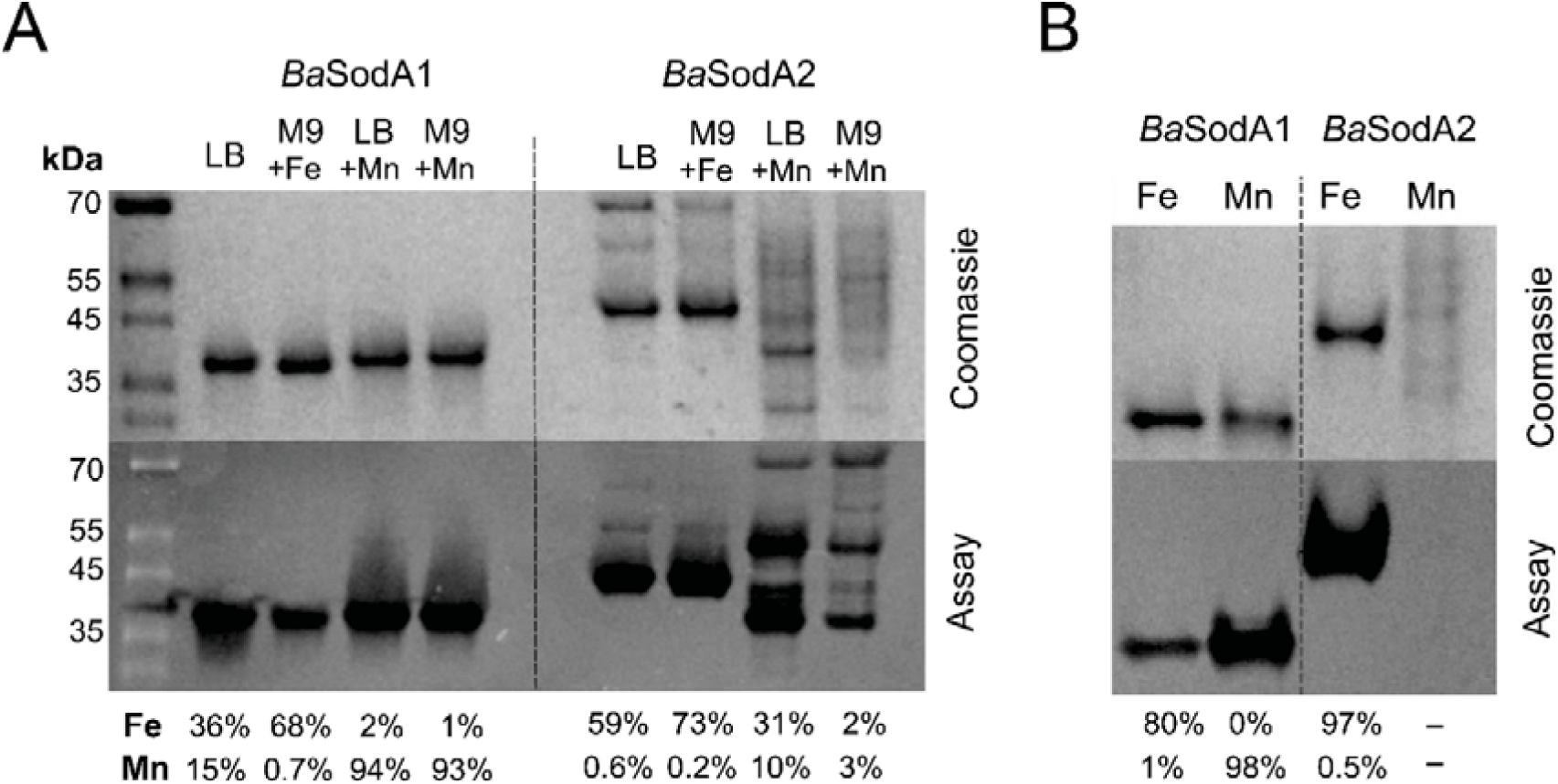
No significant Mn-incorporation by *Ba*SodA2 was achieved through metal-supplementation of expression media *or in vitro* unfolding/refolding dialysis against excess Mn. **A** Native-PAGE (upper panels) and in-gel SOD activity assays (lower panels) from analyses of purified preparations of *Ba*SodA1 (left) and *Ba*SodA2 (right) after expression either in rich (LB) or minimal (M9) media, without added metals or when supplemented with either 300 μM ammonium iron sulphate or 1 mM manganese chloride. The results from elemental analysis by ICP-MS reporting the percentage of Fe and Mn incorporation into the enzymes are annotated below. While *Ba*SodA1 could be loaded to high levels with either metal through overexpression in the presence of excess metal ion, expression of *Ba*SodA2 in Mn-supplemented media resulted in low yields of protein, which was found to be unstable, with no significant Mn-loading. Attempts to optimise expression conditions, including altering expression strain and incubation time and temperature after induction, also resulted in insignificant Mn-loading. **B** Native-PAGE and in-gel SOD activity assays from analyses of purified preparations of Fe-loaded *Ba*SodA1 and *Ba*SodA2 and of these enzymes after *in vitro* unfolding/refolding dialysis against 10 mM MnCl_2_. The refolding procedure achieved 98% Mn-loading of *Ba*SodA1, but no soluble *Ba*SodA2 remained for analysis due to total protein precipitation. In the attempt to yield soluble Mn-bound *Ba*SodA2 conditions were varied for optimisation, including protein concentration, MnCl_2_ concentration, NaCl_2_ concentration, addition of dithiothreitol to dialysis buffers and incubation time in each dialysis step. None of the conditions tested resulted in enough protein recovery for elemental analysis or spectral analysis.

**Supplementary Figure S4.**
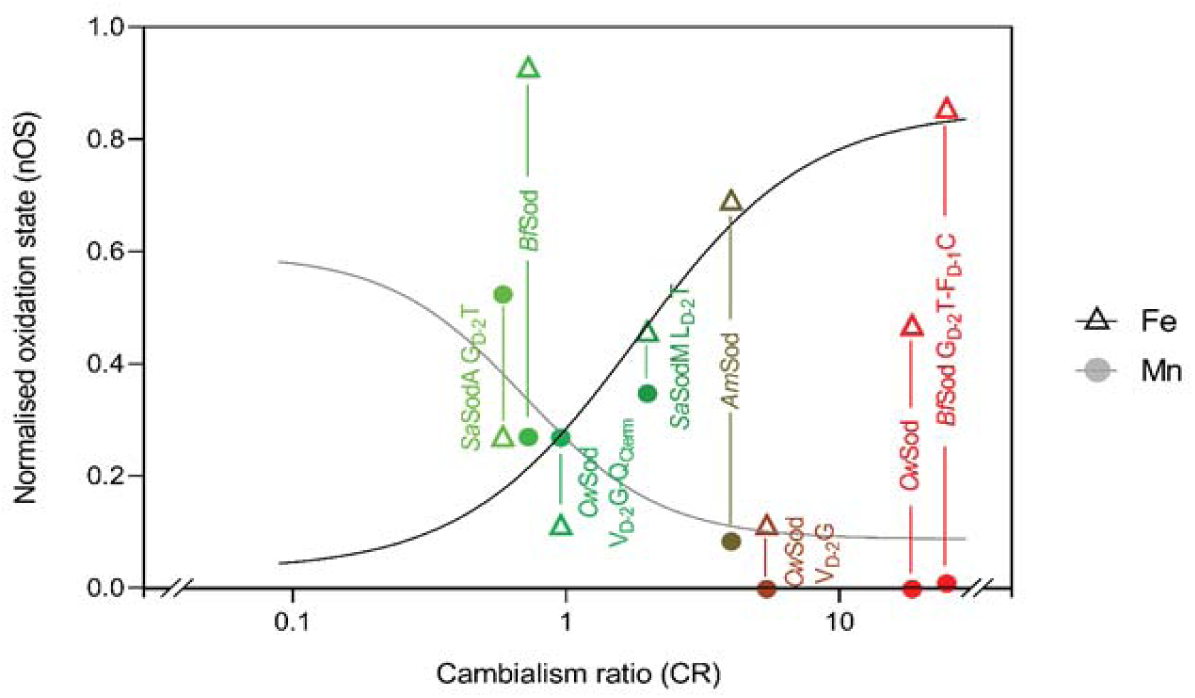
Correlation of cambialism and oxidation state extends across more divergent SODs. Normalised oxidation state (nOS) values of a further set of phylogenetically and functionally divergent SODs when Fe-loaded (open triangles) and Mn-loaded (closed circles) compared with cambialism ratio. Broadly, these SODs follow the same trend as the core dataset, despite being more evolutionarily and functionally divergent. These SODs were measured as n = 1 and were not used in the calculation of the trendline shown, which is calculated from the triplicate data shown in Figure 6. Data points are coloured by cambialism ratio and are annotated with the corresponding enzyme name.

**Table S1:** Summary of X-ray crystallography data collection statistics. Summary of parameters of the datasets collected for *Ng*SodB, *Lm*SodA and *Lm*SodA G_D-2_V-L_D-1_I protein crystals. Values in parenthesis are for the highest resolution shell.

| Data statistics |  |  |  |
| --- | --- | --- | --- |
|  | *NgSodB* | *LmSodA* | *LmSodA*<br>**G<sub>D-2</sub>V-L<sub>D-1</sub>I** |
| Beamline | I24 | I03 | I03 |
| Date | 25/02/2022 | 22/11/2021 | 22/11/2021 |
| Wavelength (Å) | 1.00 | 0.898 | 0.898 |
| Resolution (Å) | 67.47 – 2.00 (2.03<br>– 2.00) | 52.72 – 1.65 (1.68<br>– 1.65) | 51.99 – 1.40 (1.42<br>– 1.40) |
| Space group | P2<sub>1</sub>2<sub>1</sub>2<sub>1</sub> | P2<sub>1</sub>2<sub>1</sub>2<sub>1</sub> | P2<sub>1</sub>2<sub>1</sub>2<sub>1</sub> |
| **Unit-cell parameters** |  |  |  |
| a, b, c (Å) | 104.2, 105.6, 162.6 | 52.7, 94.1, 94.5 | 52.0, 93.4, 94.9 |
| α, β, γ (°) | 90.0, 90.0, 90.0 | 90.0, 90.0, 90.0 | 90.0, 90.0, 90.0 |
| Unit-cell volume (Å<sup>3</sup>) | 447110 | 117215 | 115144 |
| Solvent content (%) | 46.4 | 52.6 | 51.6 |
| Total reflections | 1596241 (79285) | 650576 (28715) | 1211451 (42183) |
| Unique reflections | 121479 (5956) | 57150 (2727) | 91660 (4487) |
| CC<sub>1/2</sub> | 1.00 (0.44) | 0.97 (0.54) | 1.00 (0.54) |
| Completeness (%) | 100 (100) | 99.7 (99.4) | 100 (100) |
| Multiplicity | 13.10 (13.30) | 11.40 (10.50) | 13.20 (9.40) |
| Rmerge | 0.16 (2.89) | 0.16 (3.17) | 0.07 (2.06) |
| I/σ(I) | 10.00 (0.90) | 7.00 (0.90) | 16.70 (1.00) |

## Author contributions

KJW, KMS and ESM conceptualised and planned the study. ESM performed the bulk of the experimental work, with contributions from KMS and RM. KMS conceptualised and performed all bioinformatic and phylogenetic analyses. AB crystallised proteins, collected and processed X-ray diffraction data and assisted EMS with the structural solution. KJW, ESM, KMS and TEKF wrote the manuscript. KJW and TEKF secured funding and managed the project.

## Acknowledgements

ESM was supported by a PhD studentship from Biotechnology and Biological Sciences Research Council (BBSRC). KMS, KJW and TEKF were supported by a grant from the National Institutes of Health (R01 AI155611) to TEKF. KJW was also funded by a Maestro grant from Narodowe Centrum Nauki (NCN), Poland (2021/42/A/NZ1/00214), which also supported RM. AB was funded by Newcastle University’s Faculty of Medical Sciences. We thank Diamond Light Source for access to beamline I03 and I24 (mx24948-124, and mx24948-132). The contents of this work are solely the responsibilities of the authors and do not reflect the official views of any of the funders, who had no role in study design, data collection, analysis, decision to publish, or preparation of the manuscript.

